# Hierarchical affinity landscape navigation through learning a shared pocket-ligand space

**DOI:** 10.1101/2025.02.17.638554

**Authors:** Bin Feng, Zijing Liu, Hao Li, Mingjun Yang, Junjie Zou, He Cao, Yu Li, Lei Zhang, Sheng Wang

## Abstract

The structure of the protein binding pocket governs the ligand binding affinity by providing crucial intermolecular interactions and spatial compatibility. While existing methods have leveraged these structural insights to advance affinity prediction, they often treat virtual screening and hit-to-lead optimization separately, mainly due to incompatible speed-accuracy requirements. However, these two tasks complement each other, and their integration enables broader chemical exploration while preserving focus on affinity-determining substructures. Here, we present LigUnity, a foundation model for affinity prediction that jointly embeds ligands and pockets into a shared space. In particular, LigUnity learns coarse-grained active/inactive distinction through scaffold discrimination and fine-grained pocket-specific ligand preference through pharmacophore ranking. We demonstrate the effectiveness and versatility of LigUnity on eight benchmarks across six settings. In virtual screening, LigUnity outperforms 24 methods with >50% improvement and demonstrates robust generalization to novel targets. In hit-to-lead optimization, it achieves state-of-the-art performance across split-by-time, split-by-scaffold, and split-by-unit settings, emerging as a cost-efficient alternative to free energy perturbation. We further showcase how LigUnity can be employed in an active learning framework for TYK2 to efficiently find optimal ligands. Collectively, these results establish LigUnity as a versatile foundation model for affinity prediction, offering broad applicability across the drug discovery pipeline.

## Introduction

The structure of the protein binding pocket, a spatially defined ligand binding cavity in proteins, serves as the foundation for computational drug design by determining non-covalent interactions and stereochemical complementarity with ligands^1^. This structure enables accurate prediction of protein-ligand binding affinity, a key determinant of drug efficacy and target selectivity. Accurately predicting this critical binding affinity plays a key role in two sequential steps in target-based drug discovery: high-throughput virtual screening, which aims to discover active compounds that can bind to an interested human protein from large-scale chemical libraries^2–4^, and hit-to-lead optimization, which aims to refine the structure of these active ligands to improve their binding affinity and drug-like properties^5,6^. For virtual screening, docking methods have been widely used with several successful applications^7^. Still, they suffer from the trade-off between the chemical search space size and computational cost^8^. As for hit-to-lead optimization, physics-based calculation methods, including free energy perturbation (FEP)^9^ and molecular mechanics-generalized born surface area (MM/GBSA)^10^, have shown encouraging performance but are either poorly correlated with experimental binding affinities or require large computational resources for extensive sampling^11,12^. To address the challenges faced by computational methods, in recent years, machine learning (ML) methods have been developed for both virtual screening (e.g., DrugCLIP^13^) and hit-to-lead optimization (e.g., ActFound^14^, PBCNet^15^). These ML approaches achieve comparable performance to computational methods while offering significant improvements in computational efficiency, thereby making them highly efficient alternatives for large-scale applications.

Despite the encouraging performance of existing ML methods, virtual screening and hit-to-lead optimization are often studied separately. However, these two tasks are interdependent and complementary to each other. A model solely focused on hit-to-lead optimization might be constrained within a limited chemical space, hindering its ability to generalize to ligands with novel chemical scaffolds. Conversely, a model developed exclusively for virtual screening might overlook crucial substructures that determine protein-ligand interactions, thus failing to distinguish between ligands with subtle structural differences. Several recent works (e.g., GenScore^16^, PIGNet2^17^, IGModel^18^, and EquiScore^19^) have begun bridging the gap between these two tasks by data augmentation techniques like re-docking, cross-docking, and decoy generation. However, these approaches still face challenges in collecting structurally similar active ligand pairs, which is crucial for learning subtle substructural variations that affect binding affinity. Based on these observations, we hypothesize that a unified foundation model jointly addressing virtual screening and hit-to-lead optimization can leverage the synergy between these two tasks to enhance overall performance.

In this work, we propose **LigUnity** (**Lig**and **U**nified affi**nity**), a protein-ligand affinity foundation model. The core innovation of LigUnity lies in its integrated capabilities for both broad screening and precise affinity prediction, achieved through a combination of scaffold discrimination and pharmacophore ranking. We jointly embed ligands and protein pockets into a shared space that captures their structural and chemical complementarity. For scaffold discrimination, LigUnity learns to differentiate between active and inactive ligands, providing insights into diverse protein-ligand interactions that help establish global structure-activity relationships across chemical scaffolds. For pharmacophore ranking, LigUnity refines the shared space by learning to order active ligands for each pocket, revealing how subtle structural differences affect binding affinity. With these computed embeddings, LigUnity enables rapid screening of large virtual libraries and efficient identification of active ligands, making it suitable for both virtual screening and hit-to-lead optimization tasks.

However, collecting a structure-aware dataset suitable for both tasks poses a significant challenge. While large-scale affinity datasets like BindingDB^20^ and ChEMBL^21^ provide abundant binding data, they lack the structural details necessary for identifying binding pocket structures. To address this, we introduce **PocketAffDB**, a comprehensive structure-aware binding assay database containing 0.8 million affinity data points across 0.5 million unique ligands and 53,406 pockets. PocketAffDB is organized by assays, in which all affinity measurements share the same experimental methods and are directly comparable. Building on this, we propose a simple but effective assay-guided pocket matching method to assign a binding pocket structure to each protein-ligand pair, based on the observation that most assays are designed for a specific binding site of interest. To our best knowledge, PocketAffDB represents the largest affinity dataset that integrates bioassay data with binding pocket structure, providing a valuable resource for affinity prediction.

To validate the effectiveness of LigUnity, we conducted comprehensive evaluations on eight benchmarks across six settings. On three virtual screening benchmarks (DUD-E^22^, Dekois^23^, and LIT-PCBA^24^), LigUnity outperforms all 24 competing methods. When applied to unseen proteins, our model achieves consistent improvement, demonstrating its robust generalization ability to novel targets. For hit-to-lead optimization, LigUnity shows superior performance on two FEP benchmarks in both zero-shot and few-shot settings, suggesting that LigUnity can be potentially used as an efficient alternative to the costly FEP calculations^9^. We further assessed LigUnity’s adaptability in split-by-time, split-by-scaffold, and split-by-unit settings on ChEMBL^21^ and BindingDB^20^ datasets, LigUnity again outperforms eight competing methods. Moreover, in an active learning framework simulating multi-iteration drug discovery optimization, LigUnity successfully identifies ligands with optimal binding affinity in just a few iterations, illustrating its practical value to minimize the experimental cost in molecular optimization. Collectively, LigUnity serves as a foundation model for both virtual screening and hit-to-lead optimization, offering broad applicability across the drug discovery pipeline.

## Results

### Overview of LigUnity

LigUnity is a protein-ligand affinity foundation model for both virtual screening and hit-to-lead optimization, employing 3D binding pocket structure to predict protein-ligand binding affinities. To train the model, we developed PocketAffDB, a comprehensive structure-aware dataset curated from large-scale experimental affinity databases (BindingDB^20^ and ChEMBL^21^) and PDB (**Fig. 1a**). We first collect affinity data into assays and then assign a binding pocket structure to each protein-ligand pair using assay-guided pocket matching (see **Methods**). To our knowledge, PocketAffDB represents the largest integrated affinity dataset combining bioassay data with structural pocket information, comprising 0.8 million affinity data points spanning 0.5 million unique ligands, 53,406 pockets, and 26,748 assays.

**Figure 1.**
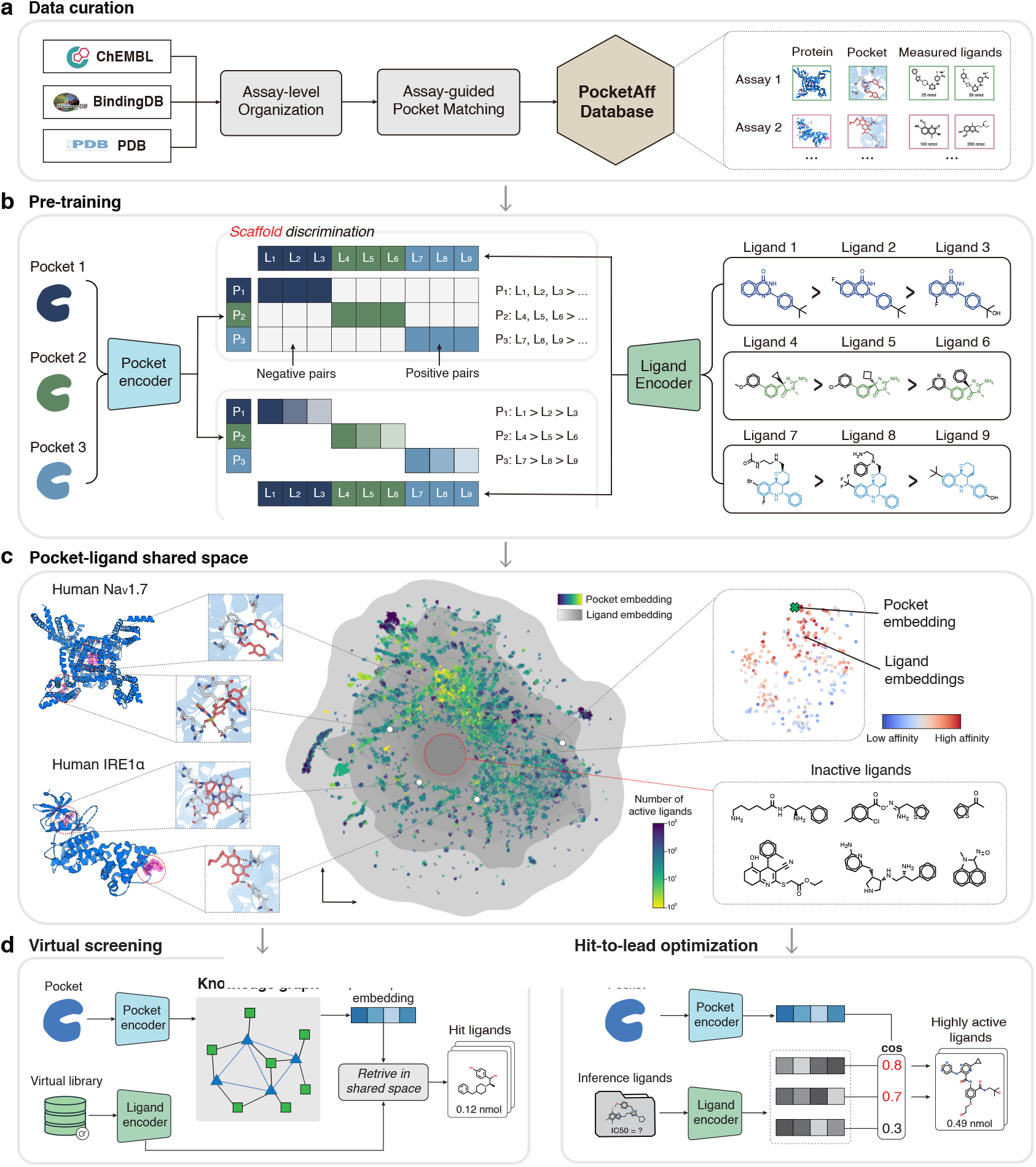
Overview of LigUnity. **a**, Data curation pipeline of PocketAffDB. **b**, LigUnity first pre-trains the pocket and ligand encoders by a hierarchical affinity landscape navigation. It first projects pockets and ligands into a shared embedding space, and then exploits scaffold discrimination to capture coarse-grained active/inactive distinction and pharmacophore ranking to refines the shared embedding space by aligning it with subtle affinity differences. **c**, Uniform manifold approximation and projection (UMAP) plot showing ligand and pocket embeddings derived by LigUnity. **d**, For virtual screening, LigUnity first refines the query pocket embedding by a graph neural network, and the updated pocket embedding is then utilized to retrieve hit ligands in the pocket-ligand shared space. **e**, For lead optimization, LigUnity is directly used to rank unmeasured ligands based on cosine similarity between pocket and ligand embeddings and find highly active ligands among them.

In the pre-training stage, LigUnity learns a pocket-ligand shared space to capture structural and chemical complementarity between pockets and ligands. We implement this through an integrated framework combining scaffold discrimination and pharmacophore ranking (**Fig. 1b**). The scaffold discrimination module learns to distinguish active/inactive compounds by focusing on structural differences in the chemical scaffold, creating an embedding space where positive pocket-ligand pairs are attracted while dissimilar negative pairs are repelled. The pharmacophore ranking component then refines this embedding space through fine-grained alignment with subtle affinity differences by focusing on the pharmacophore of ligands, where similarity rankings between ligand and pocket embeddings are used to predict relative binding affinities. These two modules complement each other, thus being employed simultaneously in the pre-training stage.

After pre-training, the embedding space of LigUnity demonstrates meaningful hierarchical patterns (**Fig. 1c**). At the coarse-grained level, ligands binding to the same pocket form visible clusters, whereas ligands targeting different pockets of the same protein are well-separated, indicating that embeddings of LigUnity can capture coarse-grained binding site information, showing potential for virtual screening. At the fine-grained ligand level, the embedding space shows that ligands with higher binding affinity values are closer to their target pockets, indicating that LigUnity learns pharmacophore-level differences affecting binding affinity, demonstrating ability for hit-to-lead optimization.

During inference, LigUnity is adapted separately for virtual screening and hit-to-lead optimization to leverage task-specific data. For virtual screening, LigUnity employs a novel graph-based approach to utilize large-scale protein-ligand interactions in existing databases (**Fig. 1d**). Specifically, we first built a large-scale knowledge pocket-ligand graph, containing 0.83 million pocket-ligand edges with known interactions and 16 million pocket-pocket edges based on structural similarity. We then applied a graph neural network to refine the query pocket embedding by aggregating neighboring information. For hit-to-lead optimization, LigUnity directly gives predictions for unmeasured ligands by simply computing cosine similarity between pocket and ligand embeddings (**Fig. 1e**). We can also enhance prediction accuracy by fine-tuning on a few experimentally measured ligands, which are often available for hit-to-lead optimization. Through modeling the affinity landscape in an embedding space, LigUnity substantially reduces the computational cost during inference, achieving six orders of magnitude speedup compared to traditional docking methods.

### LigUnity improves virtual screening

We first evaluated LigUnity for virtual screening on 102 protein targets from the DUD-E benchmark and 81 protein targets from the DEKOIS 2.0 benchmark^22,23^. These two benchmarks cover a diverse set of targets spanning enzymes, ion channels, GPCRs, and transcription factors (**Supplementary Figures. 1, 2**). We compared LigUnity with 24 different competing methods, including molecular docking methods^25,26^, structure-based methods^16,27,28^, and structure-free methods^13,29^, and observed that LigUnity outperformed all competing methods in both benchmarks (**Fig. 2a,b, Supplementary Figures. 3-6, Supplementary Table. 1**). Specifically, when compared to the best competing structure-based method Denvis-G^28^ and RTMScore^27^, LigUnity still achieves over 50% improvements in the enrichment factor (EF) 1% (*P <* 10^−9^). Meanwhile, once the embeddings are computed, LigUnity is 10^6^ times faster than Glide-SP in the screening speed as LigUnity does not need docking poses (**Supplementary Figure. 7**). To assess the ability of LigUnity to generalize to unseen proteins, we excluded training proteins that are similar to any test protein in terms of sequence similarity (**Fig. 2c,d, Supplementary Figures. 8, 9**). When using a stringent threshold of 30% sequence similarity, LigUnity still significantly outperforms both DrugCLIP and the commercial docking software Glide(SP) (P < 0.05), demonstrating its generalizability to novel proteins.

**Figure 2.**
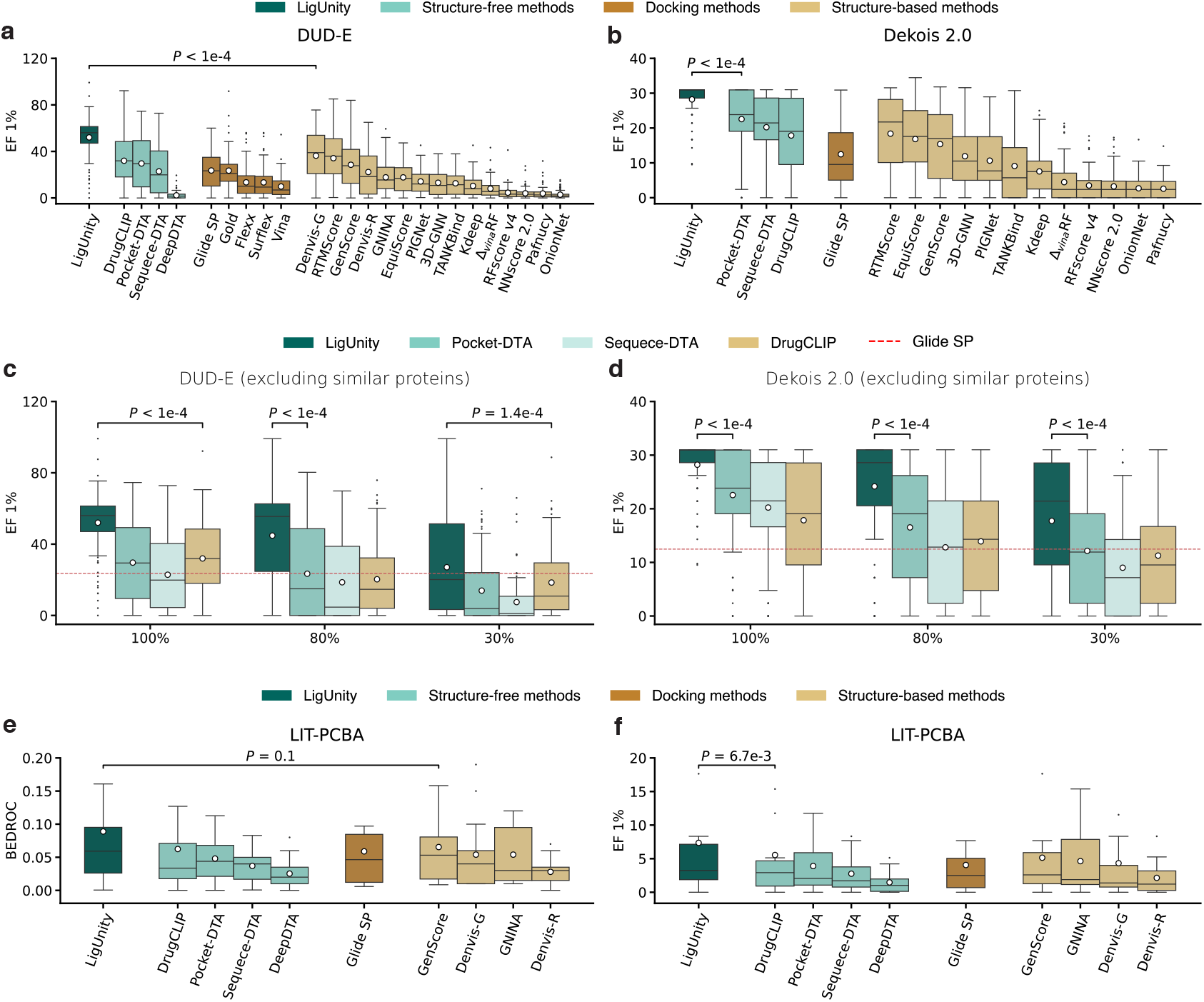
Evaluation on virtual screening. **a**,**b**, Box plots comparing LigUnity and competing methods in the DUD-E and Dekois-2.0 benchmarks in terms of enrichment factor (EF) 1%. The mean values (white dots) are calculated across n = 102 and 81 targets for the DUD-E and Dekois-2.0 benchmarks, respectively. **c**,**d**, Box plots comparing LigUnity and structure-free methods in the DUD-E and Dekois-2.0 benchmarks in terms of enrichment factor (EF) 1% using different protein training sets. The *x*-axis denotes the maximum sequence similarity between the training and test sets. The mean values (white dots) are calculated across n = 102 and 81 targets for the DUD-E and Dekois-2.0 benchmarks, respectively. **e**,**f**, Box plots comparing LigUnity and competing methods on the LIT-PCBA benchmark in terms of enrichment factor (EF) 1% and BEDROC score (*α* = 80.5). The mean values (white dots) are calculated across n = 15 targets.

Next, we evaluated LigUnity on the LIT-PCBA benchmark, which contains experimentally measured affinities for active and inactive ligands across 15 targets. Compared to DUD-E and DEKOIS, LIT-PCBA presents a more challenging setting for virtual screening, as it contains more inactive ligands that are structurally similar to active ligands. In contrast, DUD-E and DEKOIS utilize experimentally unverified decoys from the public compound database (ZINC)^30^, while excluding decoys structurally similar to active ligands to avoid potential false negatives, which may make them relatively easy for ML models. Furthermore, LIT-PCBA maintains a 1:1000 active-to-inactive ratio, which better reflects the practical conditions encountered in real-world drug discovery. Nevertheless, LigUnity still achieves superior performance on the more challenging LIT-PCBA benchmark (**Fig. 2e,f**). The improvement is also achieved when excluding training proteins that have large sequence similarity to any test protein using varying thresholds (**Supplementary Figure. 10**), reassuring the effectiveness and generalizability of LigUnity for virtual screening.

Finally, we conducted ablation studies to study the contribution of each module in LigUnity (**Supplementary Figures. 11-13**). First, we found that ablating the scaffold discrimination component leads to a performance drop of over 60% in terms of EF 1%, confirming its crucial role in distinguishing active ligands from decoys in virtual screening tasks. Second, the ablation of the heterogeneous GNN leads to consistent decreases in performance, demonstrating its capability to utilize information from the pocket-ligand heterogeneous graph to improve the predictive power. Furthermore, we found that the pharmacophore ranking is particularly effective on LIT-PCBA among three benchmarks (**Supplementary Figure. 13**), as it helps LigUnity distinguish subtle structural differences among similar ligands that are crucial to their affinity scores. Together, our ablation studies demonstrate the importance of all three key technical ideas proposed by LigUnity, necessitating the joint optimization of virtual screening and hit-to-lead optimization.

### LigUnity improves hit-to-lead optimization

Hit-to-lead optimization aims to optimize the structure of the ligand for improved affinity by ranking candidate ligands that are commonly structurally similar to initial hits identified to be active. After observing the substantial improvement of LigUnity in virtual screening, especially its ability to capture ligand structure differences, we next investigated whether LigUnity can be applied to hit-to-lead optimization. We evaluated LigUnity on two FEP benchmarks (JACS^9^ and Merck^31^) to predict binding free energies, which are the most widely used indicators for hit-to-lead optimization. These two benchmarks contain 16 targets, and each target has an average of 29 experimentally measured ligands. Following previous work^14^, assays similar to those included in the FEP benchmarks are excluded in the pre-training stage to avoid potential data leakage (see **Methods**). Ligands in the FEP benchmarks are also removed from the pre-training data.

In the zero-shot setting, where the model does not use any measured ligands for test proteins during pre-training and fine-tuning, our model surpasses existing computational methods (e.g. Glide-SP^25^, MM/GBSA^32^), structure-based methods (e.g. GenScore^16^), and structure-free methods (e.g. DrugCLIP^13^) (**Fig. 3a**), on 8 targets of the Merck FEP benchmark, demonstrating the effectiveness of our method. To further examine the performance of LigUnity in hit-to-lead optimization, we studied three more challenging settings: (1) no similar ligands setting, where training ligands with more than 50% Tanimoto similarity to any test ligand are excluded; (2) no similar proteins setting, where training proteins with more than 40% sequence similarity to any test protein are excluded^33^; (3) no similar ligands and proteins setting, where the criteria of the first two settings are both applied to exclude similar proteins or ligands. LigUnity again consistently outperforms competing methods across all three settings (**Supplementary Figure. 14**). Notably, on 16 targets from Merck and JACS benchmarks, it achieves a significant improvement of 38.1% over the sequence-only variant LigUnity(seq) under the most difficult no similar ligands and proteins setting (15.4% under the no same ligands setting), demonstrating the importance of incorporating pocket information for better generalization to novel proteins.

**Figure 3.**
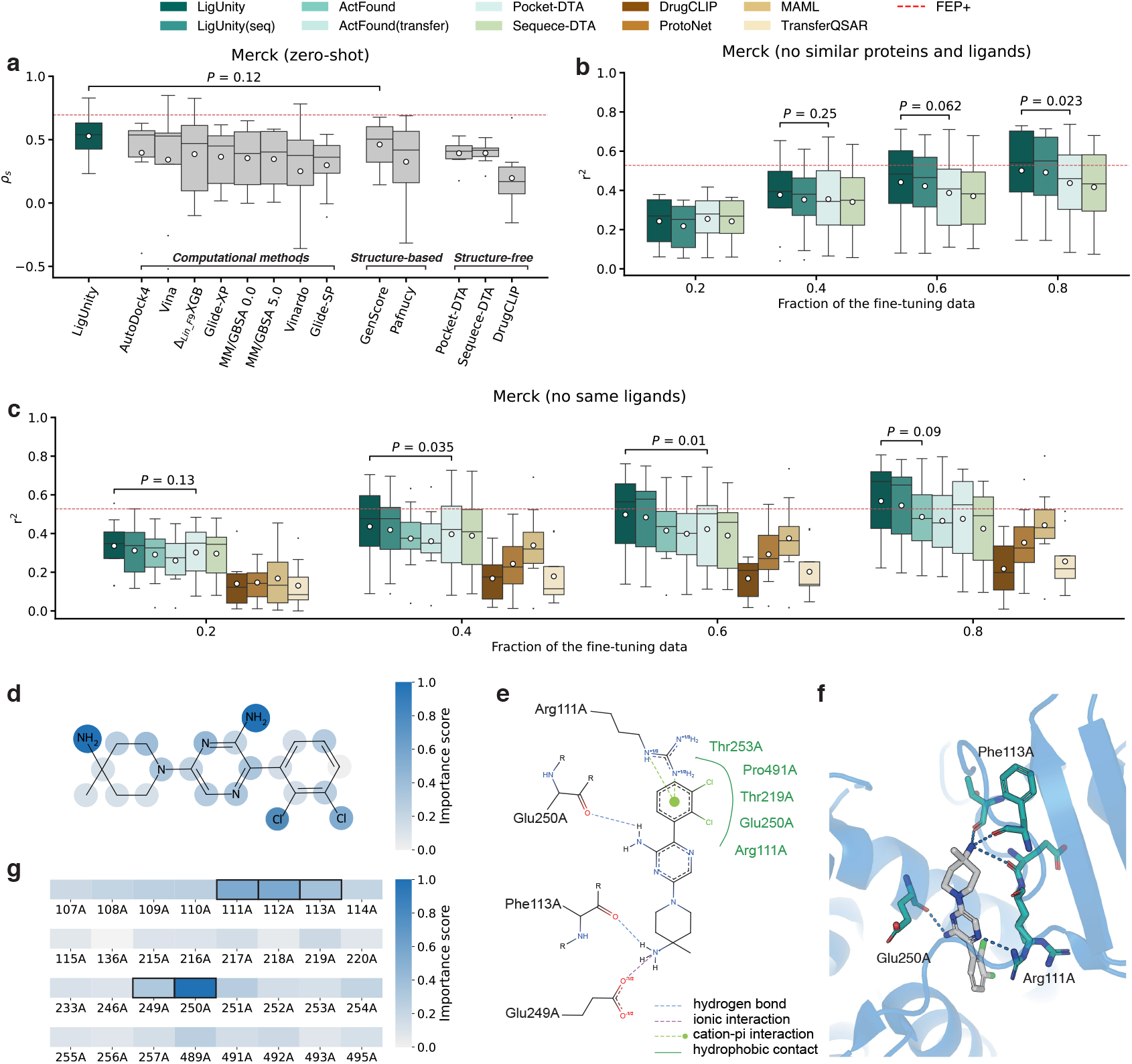
Evaluation on hit-to-lead optimization. **a**, Box plot comparing the binding affinity prediction on the Merck benchmark in terms of Spearman’s rank correlation (*ρ*_*s*_) in the zero-shot setting. The mean values (white dots) are calculated across n = 8 targets. **b**,**c**, Box plots comparing the binding affinity prediction across different settings on the Merck benchmark in terms of *r*^2^ when 20%, 40%, 60%, 80% of the experimental binding affinities are used for fine-tuning. The mean values (white dots) are calculated across n = 8 targets. **d**, Case study on the SHP2 target showing the importance score of each ligand atom predicted by LigUnity. **e**,**f**, 2D **(g)** and 3D interaction graph **(h)** showing the non-covalent interaction between the ligand and pocket (PDB ID: 5EHR). **g**, Case study on the SHP2 target showing the importance score of each pocket residue predicted by LigUnity. The top five predicted key residues are highlighted with black boxes.

In real-world drug discovery, binding data for a few ligands of interest are often experimentally measured. A successful hit-to-lead optimization model should be able to leverage these binding data to enhance its predictions. To examine the ability to employ these data, we fine-tuned LigUnity and competing methods with varying proportions of ligand binding data. We observed that LigUnity consistently outperforms competing methods across all four settings (**Fig. 3b,c, Supplementary Figure. 15, Supplementary Table. 2**). Even under the most challenging setting where both similar ligands and proteins are excluded, LigUnity achieves an *r*^2^ of 0.501 on the Merck benchmark after fine-tuning on 80% of ligands (23.2 ligands on average for each target), reaching performance comparable to FEP+(OPLS4) (*r*^2^ = 0.528), a leading commercial computational software for calculating the binding free energy. These results demonstrate the practical value of LigUnity for affinity prediction, offering an accurate and efficient alternative to resource-intensive computational chemistry approaches.

Similar to our analyses in the virtual screening, we conducted ablation studies to investigate the contribution of each component in LigUnity (**Supplementary Figure. 16**). Our results demonstrate that excluding the scaffold discrimination or the pharmacophore ranking component leads to substantial performance drops. Notably, ablating the pharmacophore ranking leads to a performance drop of over 50% in both zero-shot and few-shot settings, highlighting the effectiveness of the ranking objective in capturing the relative activity differences for hit-to-lead optimization tasks. To verify that LigUnity captures meaningful interaction patterns, we systematically tested it on FEP benchmarks by masking interaction-critical residues. Key residues were identified from PDB structures using ProteinPlus^34^. The results show that masking strong H-bond/ionic bond residues caused significant performance drops (Δ*ρ*_*s*_ = −6.34%), versus minimal impact from random masking (Δ*ρ*_*s*_ = −0.46%) (**Supplementary Table. 3**), demonstrating that LigUnity’s ability to rely on protein-ligand interactions to achieve good performance.

Finally, to help understand the structural significance of individual atoms and residues involved in the interaction, we calculated the importance score of each atom and protein pocket residue on the SHP2 target by masking them individually (**Fig. 3d-g, Supplementary Figure. 17**, see **Methods**). By examining the atom importance scores, LigUnity correctly identifies two amino groups (-NH2) as key contributing factors to the predicted binding affinity (**Fig. 3d**), aligning with the crystallographic observations of their hydrogen bonding and ionic interactions (**Fig. 3e,f**). Furthermore, using protein residue importance scores, the model correctly highlights the crucial role of Glu250E in hydrogen bonding, along with the polar interactions involving Phe113A, Arg249A, and Arg111A (**Fig. 3g**). Although the pose-free nature of our approach results in certain limitations, such as underestimating the contributions from specific hydrophobic cavity residues (Thr253A, Pro491A), LigUnity successfully identifies key interaction patterns. These results suggest that LigUnity can serve as an interpretable computational tool for studying structure-activity relationships and guiding hit-to-lead optimization.

### LigUnity as a versatile foundation model for different applications

To further explore the ability of LigUnity as a foundation model for affinity prediction, we next sought to evaluate its performance on assays from ChEMBL^21^ and BindingDB^20^, two comprehensive public databases containing experimentally measured binding affinities covering various aspects of assay types, such as protein-based, cell-based, cell-membrane-based, and subcellular-based assays^20,21^. We first used a split-by-time setting to evaluate the performance, where the model is pre-trained using 18,552 assays released before March 2019 and tested on 161 assays released after March 2019 in ChEMBL and BindingDB. Each test assay has 48.6 ligands on average. To avoid potential data leakage, we further excluded test assays whose targets exist in the pre-training data.

LigUnity attains consistent improvement using varying numbers of fine-tuning data ranging from 4 to 16 (**Fig. 4a, Supplementary Figure. 18**). When compared to ActFound^14^, the best-competing method specifically designed for few-shot affinity prediction, LigUnity achieved an average improvement of 8.5% in terms of *r*^2^. Since ActFound is a ligand-based model without considering pocket information, our substantial improvement reflects the effectiveness of using protein pocket information for affinity prediction. On a more challenging split-by-scaffold setting where test ligands have distinct scaffolds from training ligands, LigUnity again obtains the best performance (**Fig. 4b**), demonstrating its strong generalizability to unseen chemical scaffolds.

**Figure 4.**
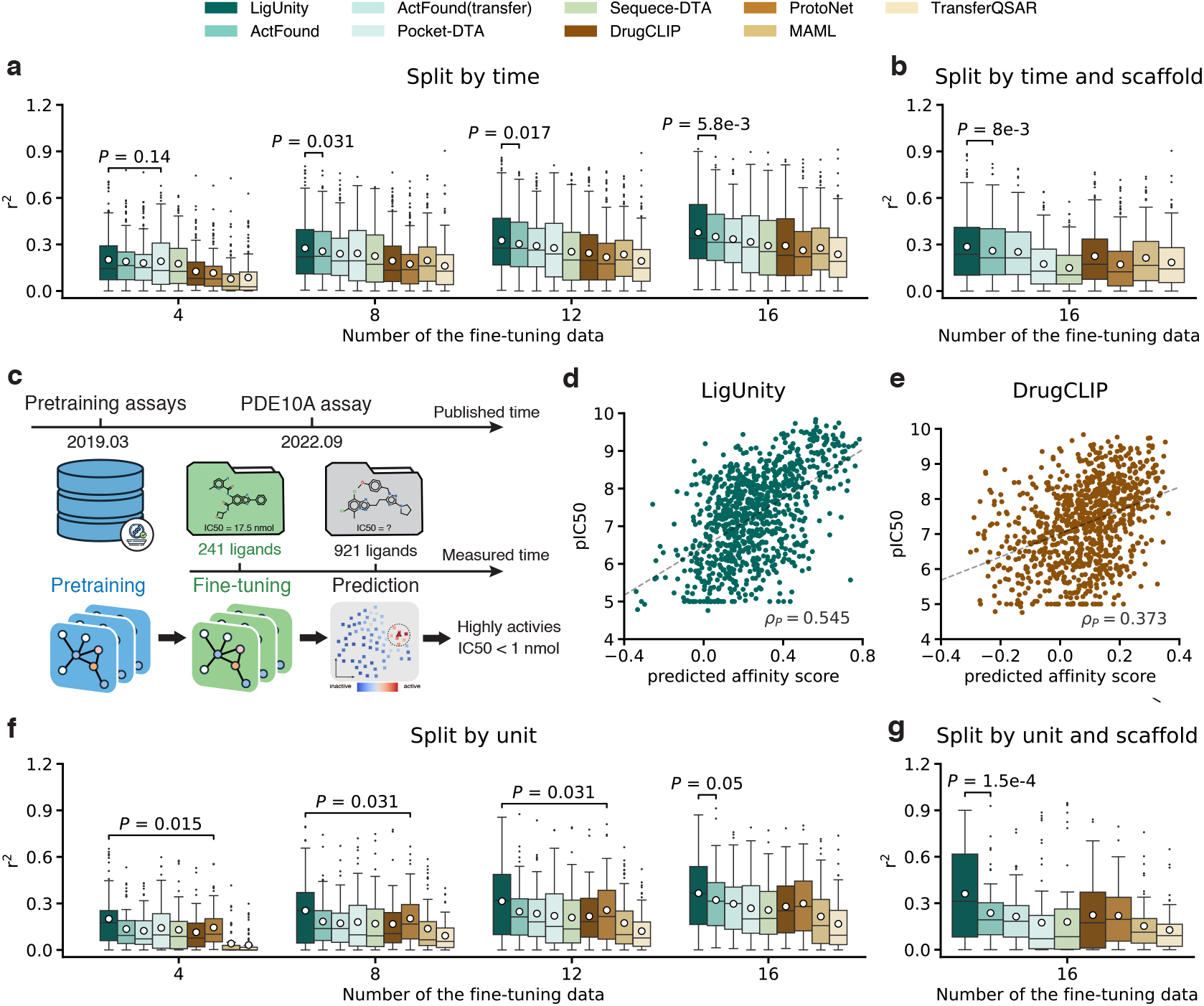
Evaluation on diverse settings. **a**,**b**, Box plots comparing the binding affinity prediction on the split-by-time setting in terms of *r*^2^ when 4, 8, 12, 16 experimentally measured ligands are used for fine-tuning. The mean values (white dots) are calculated across n = 161 assays. **c**, Experimental setting for case study on the phosphodiesterase 10A (PDE10A) target. **d**,**e**, Scatter plots comparing the predicted affinity score and pIC50= −log_10_IC50 for LigUnity (**f**) and DrugCLIP (**g**) when finetuned with the first measured 20% ligands. **f**,**g**, Box plots comparing the binding affinity prediction on the split-by-unit setting in terms of *r*^2^ when 4, 8, 12, 16 experimentally measured ligands are used for fine-tuning. The mean values (white dots) are calculated across n = 65 assays.

Next, we conducted a case study on phosphodiesterase 10A (PDE10A), a verified therapeutic target for antipsychotic therapy with no approved drugs, to assess the applicability of LigUnity in the real-world drug discovery pipeline. This dataset was released by Roche Ltd. in 2022, comprising 1,162 ligands with experimentally measured binding affinity and timestamp of each experiment^35^. We used a split-by-time setting for ligands in the PDE10A dataset, where the first measured 20% ligands are used for fine-tuning and the remaining 80% are used for testing (**Fig. 4c**). To ensure a rigorous split-by-time evaluation, we also restricted the pre-training data to the assays released before March 2019. To this end, our fine-tuned model achieves a Pearson correlation of 0.55 on the test set (**Fig. 4d**), substantially outperforming the 0.37 Pearson correlation of DrugCLIP (**Fig. 4e**). Building on this performance, our model demonstrates a strong ability to identify highly potent ligands, successfully identifying 4 highly active ligands (IC50 *<* 1 nmol) among the top-10 predicted ligands (**Supplementary Figure. 19**) and 24 highly active ligands among the top-50 predicted ligands (compared to 11 for DrugCLIP). Collectively, these results demonstrate the superior capability of LigUnity in identifying highly potent ligands, holding the potential to reduce the cost of hit-to-lead optimization.

Finally, we studied a split-by-unit setting where the test assays use percentage (“%”) units, a widely used format (24.4% in ChEMBL) in practical drug design with limited exploration in ML methods. Assays with percentage units pose challenges for affinity prediction, as they distribute differently from pre-training assays with molar concentration (e.g., nmol) or density units (e.g., *µ*g*/*ml). We hypothesize that our pharmacophore ranking approach enables LigUnity to be robust against assays with different units, as it aims to learn the pocket-specific ligand ranking instead of the absolute affinity value. To this end, we examined a split-by-unit setting by using ChEMBL assays reporting activities in percentage (%) as test assays and assays with defined molar concentration or density units as pre-training assays. This test set contains 65 assays, and each assay has on average 30.5 ligands. When fine-tuned with different numbers of ligands (4 to 16), LigUnity consistently surpasses all competing methods (**Fig. 4f,g, Supplementary Figure. 18**). Notably, compared to the regression model Pocket-DTA, LigUnity shows a 40.2% improvement in this split-by-unit setting, substantially higher than the 13.8% improvement in the split-by-time setting, demonstrating the generalization ability of the pharmacophore ranking approach across different measurement units, reassuring LigUnity as an effective foundation model for drug discovery.

### LigUnity boosts the active learning framework for drug discovery

In the real-world drug discovery pipeline, multiple iterations of molecular optimization are often required due to limited experimental resources and the difficulty in accurately predicting protein-ligand binding affinity. As a result, active learning is often exploited in this process. Building on this, we explored how LigUnity can be integrated with active learning to optimize ligands for Tyrosine Kinase 2 (TYK2), a therapeutic target for autoimmune diseases. We employed a dataset that contains binding free energy for 10,000 ligands calculated using FEP, consuming approximately 80,000 GPU hours (9.1 GPU years)^36^. Following established active learning protocols^37^, we aimed to identify highly active ligands with limited computational resources in terms of the amount of FEP calculations.

To this end, we integrated LigUnity in an active learning framework, where we first trained the model using a small number of randomly selected ligands with known binding free energies. In each subsequent iteration, we selected a subset of unlabeled ligands and performed FEP calculations to determine their binding free energies. These newly labeled data points are then incorporated into the training set, and the model is retrained to refine its predictions (**Fig. 5a**).

**Figure 5.**
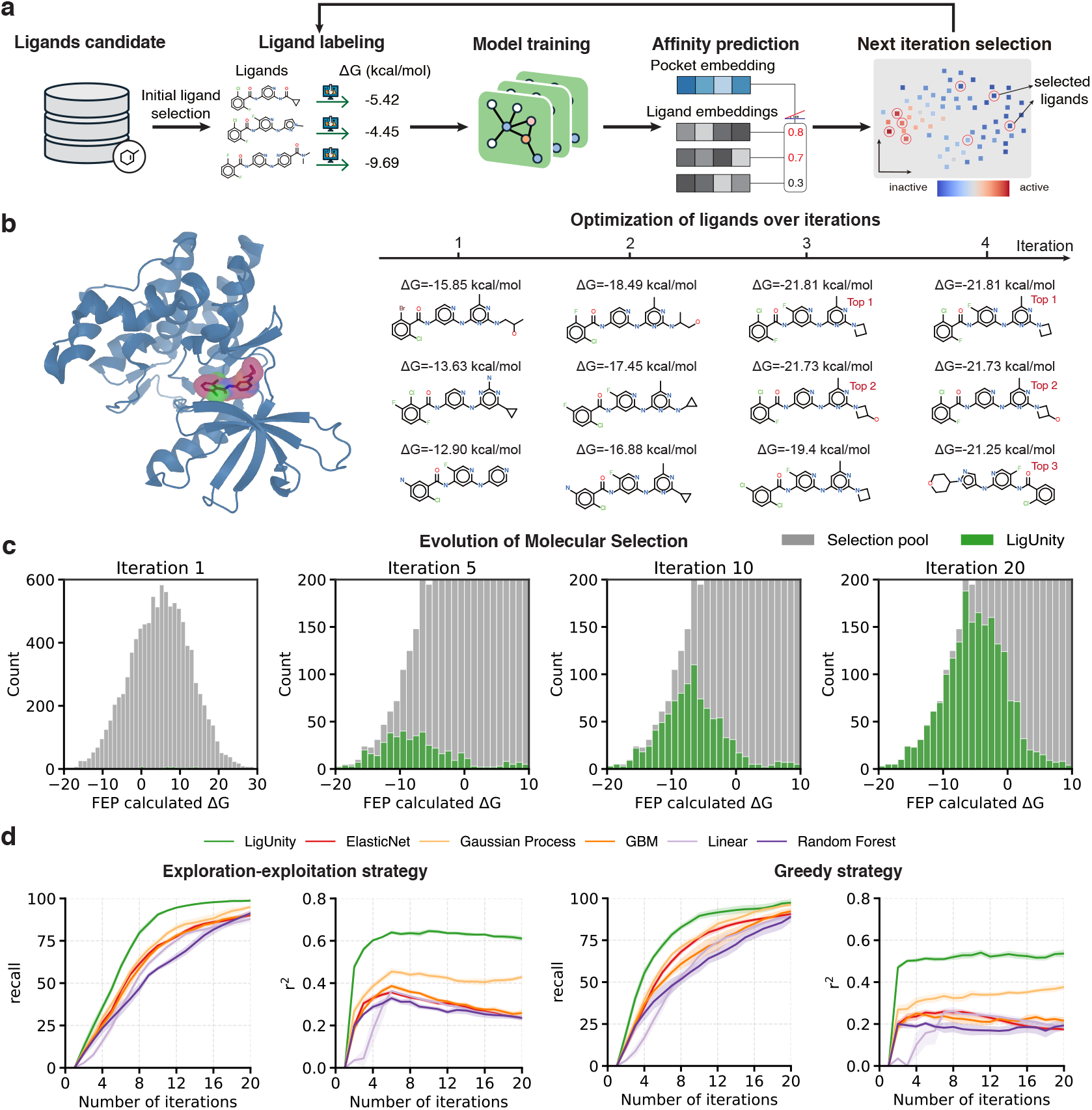
Evaluation of LigUnity in an active learning framework on TYK2. **a**, An active learning framework based on LigUnity. **b**, 3D structure for the TYK2 target (left panel) and the top ligands selected for iteration 1, 2, 3, and 4 (right panel) under the exploration-exploitation strategy, FEP-calculated binding free energies for each ligand are also shown. **c**, Histogram plots showing the distribution of ΔG values of selected ligand for each iteration based on a greedy selection strategy. **d**, Plots comparing LigUnity and competing methods on the TYK2 dataset in terms of top 2% recall and *r*^2^ when using the exploration-exploitation strategy (left) and greedy selection strategy (right).

We compared all methods under two selection strategies: a greedy strategy and an exploration-exploitation strategy (see **Methods**). We first found that LigUnity successfully identified best active ligands within a few iterations (**Fig. 5b**). The proportion of highly active ligands continually increased after each iteration (**Fig. 5c**), demonstrating the effectiveness of the active learning framework in identifying highly active ligands. Moreover, we found that LigUnity consistently outperforms other competing methods under both greedy and exploration-exploitation strategies, achieving over 40% improvement in terms of *r*^2^ (**Fig. 5d**). By comparing two different selection strategies, we found that the greedy strategy has a higher recall in early iterations by focusing on the most promising candidates (**Fig. 5d, Supplementary Figure. 20**). Interestingly, the exploration-exploitation strategy attains superior performance in later iterations, primarily because of its more diverse ligand sampling in early iterations that helps improve model accuracy. These findings demonstrate the broad applicability of LigUnity in drug design, particularly in identifying highly active ligands using limited experimental and computational resources.

## Discussion

We have presented LigUnity, a protein-ligand affinity foundation model for both virtual screening and hit-to-lead optimization. Our model learns a pocket-ligand shared space to capture structural and chemical complementarity between pockets and ligands, which is implemented through an integrated framework combining scaffold discrimination and pharmacophore ranking. For virtual screening, our experiments have shown that LigUnity outperformed 24 competing methods across three benchmarks (DUD-E, LIT-PCBA, and Dekois 2.0). For hit-to-lead optimization, our model achieves the best performance in two FEP benchmarks and demonstrates promising accuracy in split-by-time, split-by-scaffold, and split-by-unit settings. Moreover, by developing an active learning framework to simulate real-world lead optimization, LigUnity efficiently identifies optimal binding compounds with limited computational resources. Collectively, these results demonstrate the strong generalization ability and practical usability of LigUnity throughout the early-stage drug discovery pipeline, establishing our confidence in LigUnity as a versatile foundation model for computer-aided drug discovery.

Two main lines of work are related to LigUnity, including structure-free Drug-Target Affinity (DTA) prediction methods and structure-based methods. Compared to DTA methods^29,38–46^, which directly predict binding affinity using protein and ligand information, LigUnity presents three key differences. First, DTA methods are challenged by the small number of proteins with measured affinity (2,773 in BindingDB), whereas LigUnity incorporates fine-grained pocket information and leverages a comprehensive training set covering 53,406 pockets for 4,847 proteins, enabling robust performance on under-studied proteins. Second, existing DTA methods perform direct regression of absolute affinity values, whereas LigUnity learns a pocket-ligand shared space using scaffold discrimination and pharmacophore ranking, reflected by the consistent performance improvement in our experiments. Third, DTA methods require intensive computation for each prediction, whereas LigUnity enables rapid screening by pre-computing and storing embeddings, leading to better efficiency for large-scale applications.

Compared to structure-based methods^16,19,27,28,47–57^that rely on 3D binding poses, LigUnity presents two improvements: First, structure-based methods are constrained by the scarcity of ligands with co-crystal structures. For example, only 7.5% of 0.57 million compounds with measured activities in BindingDB have crystal structures^20^. In contrast, LigUnity is developed using a diverse training set of 0.45 million unique ligands, enabling effective generalization to novel chemical scaffolds; Second, structure-based methods require computationally expensive pose generation (approximately one minute per compound using Glide), whereas LigUnity achieves competitive performance without the need for pose information, making it applicable to large-scale screening. Furthermore, our approach offers significant flexibility for diverse scenarios. For novel targets with limited structural data, LigUnity can leverage predicted pockets from AlphaFold 3 or homology modeling. In cases where no experimental or predicted structures exist, we have also provided sequence-based variants of LigUnity.

Despite its promising performance in binding affinity prediction, LigUnity has two limitations that can be addressed in future works. First, LigUnity can only be applied to assays with known protein targets, limiting its applications to diverse assays without target information (e.g., phenotype assays). To expand the applicability of our model, we plan to incorporate additional modalities such as gene information in future work. Second, although our current approach achieves strong results without structural information, integrating both binding pose data and activity data could potentially enhance prediction accuracy, particularly in scenarios where sufficient computational resources are available to obtain precise binding poses.

## Method

### Problem setting

LigUnity aims to predict binding affinity between given binding pockets and unmeasured ligands. Our training data are organized by assay, which is defined as the experimental procedure designed to evaluate the binding affinity of ligands against a specific protein target. Each assay has multiple measured ligands and only one target protein. A key consideration is that affinity values are only comparable within the same assay, and comparing affinity values from different assays targeting the same protein may not be reliable^58^. This incomparability arises from variations in experimental conditions (e.g., cofactor concentration, pH, temperature), assay formats (e.g., cell-based vs. target-based assays, different detection methods), and affinity measurements (e.g., IC50, Ki, Kd). Therefore, our method focuses on learning relative affinity rankings within each assay instead of absolute values.

Let 𝒜 denote the set of assays. For each assay *A*_*i*_ *∈* 𝒜, *L*_*i*_ denotes the set of tested ligands, and *v*_*i*_(*l*) denotes the affinity value of ligand *l ∈ L*_*i*_. Each assay corresponds to one target protein, which may have multiple PDB structures available. We define *P*_*i*_ as the set of protein pocket structures for assay *A*_*i*_ and randomly sample one pocket structure from *P*_*i*_ at each training step.

### Pocket and ligand encoder

In the pre-training stage, our pocket encoder and ligand encoder are initialized with Uni-Mol^59^, an SE(3)-invariant graph transformer for 3D moleculars, which is a structure-free method and does not rely on the binding pose as input. For the ligand branch, we first use RDKit to generate 3D conformations for each ligand which have been proven effective for various tasks^60^, and then input them into the Uni-Mol ligand encoder to generate ligand embeddings. Similarly, for the pocket branch, we input the protein pocket conformation into the Uni-Mol pocket encoder to generate the protein embedding. For a pocket-ligand pair (*p, l*), their vector representations are obtained through:

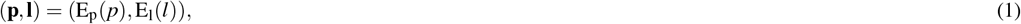

where E_p_ and E_l_ denote the pocket and ligand encoders respectively, **p** denotes the protein embedding, and **l** denotes the ligand embedding.

### Hierarchical affinity landscape navigation

In the pre-training stage, LigUnity learns a pocket-ligand shared space using two complementary strategies: scaffold discrimination and pharmacophore ranking. The scaffold discrimination approach captures the coarse-grained distinction between active and inactive pocket-ligand pairs by focusing on structural differences in the chemical scaffold, which aims to bring the embeddings of positive pocket-ligand pairs closer and push the negative pairs apart. The pharmacophore ranking approach learns to align the embedding space with the fine-grained affinity ranking of measured ligands by focusing on the pharmacophore. These two approaches complement each other, and we jointly optimize them to make the model suitable for both virtual screening and hit-to-lead optimization scenarios.

During inference, the model only needs to take 3D protein pockets and 3D ligand conformations (generated by RDKit) as input. It then calculates the pocket and ligand embeddings and outputs their cosine similarity as the predicted affinity score^13^. This differs from most previous machine learning affinity prediction models, which typically model affinity prediction as a regression task and use either dual-tower architectures (structure-free methods)^29,42,44^ or require binding poses (structure-based methods)^27,28^ as input. Once the pocket and ligand embeddings are pre-computed, LigUnity enables fast screening by similarity-based comparison in joint embedding space, achieving six magnitudes of speedup compared to docking methods.^13^

### Pharmacophore ranking

Pharmacophore ranking is an approach that prioritizes active compounds based on their binding potential and structural nuances. We implement pharmacophore ranking through listwise learning. Unlike pointwise approaches that predict absolute scores or pairwise approaches that compare item pairs, listwise ranking considers the entire ranked list simultaneously, making it particularly suitable for problems where the relative ordering matters more than absolute values. This approach allows us to capture subtle structural differences between ligands that share similar scaffolds but exhibit varying binding affinities to the same pocket. We adopt the Plackett-Luce model^61^, a statistical model specifically designed for ranking data. The fundamental idea of the Plackett-Luce model is to assign each item a positive “strength” parameter, which determines the probability of the item being selected. In our context, the strength parameter *θ*_*i*_ for each ligand is computed as the exponential of its pocket-ligand similarity score: *θ*_*i*_ = exp(sim(**p, l**_*i*_)). For a set of items with strength parameters {*θ*_1_, …, *θ*_*m*_}, the probability of selecting item *i* is defined as:

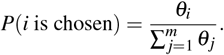

The Plackett-Luce model treats ranking as a sequential selection process. Given *n* ligands to rank, the model first selects the top-ranked ligands from all candidates using the above probability formula. Then, it selects the second-ranked ligand from the remaining *n* − 1 candidates, again using the same probability formula but only considering the remaining ligands. This process continues until all ligands are ranked. The probability of observing a complete ranking sequence *π* is the product of these conditional probabilities at each selection step:

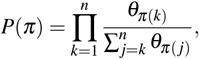

where *π*(*k*) denotes the index of the ligand at position *k* in the ranking. The model parameters are optimized by minimizing the negative log-likelihood of observing the true ranking based on measured affinity values. This framework naturally handles varying numbers of ligands across different assays and provides a probabilistic interpretation of the ranking process.

Given this probabilistic framework, we can define the ranking loss for assay *A*_*i*_ as the weighted negative log-likelihood of the observed ranking probability. For each assay *A*_*i*_ *∈* 𝒜, we order ligands according to their measured affinity values. At the *k*-th step of the selection process, considering all ligands with affinity values no greater than that of the *k*-th ranked ligand, we define the selection probability for the *k*-th ranked ligand as:

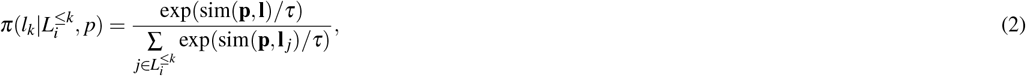

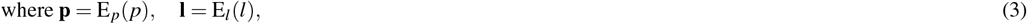

where *p ∈ P*_*i*_ is the 3D structure of the pocket tested in the assay *A*_*i*_; *l*_*k*_ the *k*-th ranked ligand; 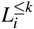 represents the set of ligands with affinity values no greater than *v*_*i*_(*l*_*k*_); *τ* is a temperature hyper-parameter; and sim(**p, l**) denotes the cosine similarity between protein and ligand embedding vectors.

Based on these step-wise selection probabilities, we compute the ranking loss as the weighted sum of the negative log probabilities over all selection steps:

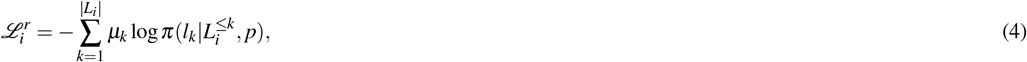

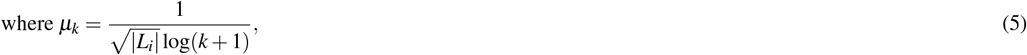

where *µ*_*k*_ is a decay factor with two components: 1*/* log(*k* + 1) prioritizes top-ranked molecules and 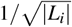 prevents assays with many ligands from dominating the training loss. We also experimented with 1*/* |*L*_*i*_| and constant scaling terms and empirically found that 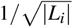 yields the best performance.

### Scaffold discrimination

Scaffold discrimination, implemented through contrastive learning, aims to maximize the similarity between active pocket-ligand pairs while minimizing the similarity between inactive pairs. This approach enables the model to learn global structure-activity relationships across diverse chemical scaffolds, providing a foundation for broadly distinguishing active from inactive compounds. Following Gao et al.^13^, we formulate our contrastive loss using in-batch softmax, which consists of two components: the pocket-to-ligand loss 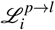 and ligand-to-pocket loss 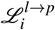:

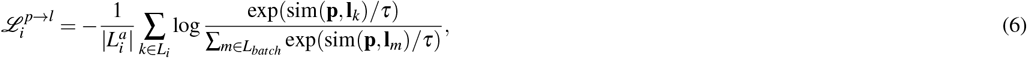

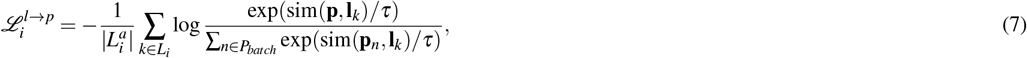

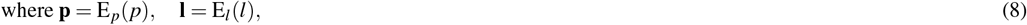

where *p ∈ P*_*i*_ is a 3D structure of the pocket tested in the assay *A*_*i*_; *L*_*batch*_ and *P*_*batch*_ denote the set of ligands and pockets sampled in this training batch, respectively; *τ* is a temperature hyper-parameter, and 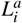 denotes the set of active ligands tested in the assay *A*_*i*_.

Building on this formulation, we define how positive and negative pairs are identified in our framework: pocket-ligand pairs with experimentally measured affinity (*≤*10*µ*mol) are treated as positive pairs following Mayr et al.^62^, while pairs without measured affinity values serve as negative pairs. Although the boundary between active and inactive ligands varies across different protein targets, our framework naturally addresses this issue by learning the relative affinity among ligands rather than relying solely on binary active/inactive labels. However, this in-batch negative sampling may introduce potentially harmful false negative pocket-ligand pairs. To mitigate this, we devised a simple strategy to identify potentially active pocket-ligand pairs. Specifically, pocket-ligand pairs formed between assay A and assay B are excluded from negative sampling if the protein targets of the two assays share the same UniProt ID. This exclusion is based on the observation that protein pockets with the same UniProt ID are likely to share overlapping active ligands.

The total contrastive loss is defined as:

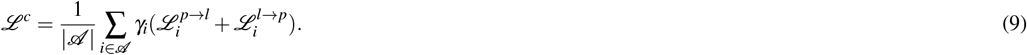

where 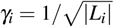 prevents assays with many ligands from dominating the training loss.

### Pre-training objective

To optimize both virtual screening and hit-to-lead optimization, LigUnity is pre-trained with a combined loss that integrates scaffold discrimination and pharmacophore ranking:

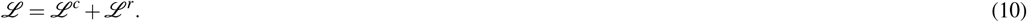

Collectively, this hierarchical approach enables LigUnity to navigate the affinity landscape by first broadly discriminating active scaffolds from inactive ones, and then precisely ranking ligands based on their pharmacophoric features, supporting its effectiveness in both virtual screening and hit-to-lead optimization tasks. We demonstrated that jointly optimizing these two tasks can help each other, which is shown in our ablation studies (**Supplementary Figures. 13, 16**).

### Heterogeneous GNN

Based on the observation that structurally similar pockets tend to have similar interaction patterns with ligands, we hypothesize that integrating information from existing large-scale pocket-ligand databases should improve the performance of LigUnity. To this end, we propose a Heterogeneous Graph Neural Network model (H-GNN) to refine pocket embeddings by incorporating information from a large-scale pocket-ligand knowledge graph.

Our H-GNN model consists of two main components: a pocket-pocket aggregator and a pocket-ligand aggregator. This model receives a directed knowledge graph *G* = (*V*_*p*_,*V*_*l*_, *E*_*l→p*_, *E*_*p→p*_) of pockets and ligands as input, where *V*_*p*_ = {**p**_*i*_},*V*_*l*_ = {**l**_*i*_} are two sets of nodes representing the pockets and ligands respectively; *E*_*l→p*_ is the ligand-to-pocket edge set and *E*_*p→p*_ is the pocket-to-pocket edge set. For ligand-to-pocket edges, we add an edge from a ligand to a pocket if the ligand is active for the protein pocket with experimentally measured affinity. For pocket-to-pocket edges, we add edges between pockets based on sequence alignment scores, which are computed using scikit-bio with the BLOSUM50 substitution matrix. We only add edges between pockets with high alignment scores. Specifically, if the alignment score between pocket A and pocket B is larger than half of the self-alignment score of pocket A, we add an edge from pocket B to pocket A.

We constructed this large-scale pocket-ligand knowledge graph from our pre-training data. The graph contains 53,406 pocket nodes, 0.45 million ligand nodes, 0.83 million ligand-to-pocket edges, and 16 million pocket-to-pocket edges. During the virtual screening, the H-GNN model refines the embedding of the query protein pocket by incorporating information from the heterogeneous graph: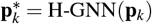. We then use this refined pocket embedding 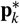 to retrieve hit ligands in the candidate ligand embedding space.

### Pocket-pocket aggregator

We first use pocket-pocket aggregator to incorporate information from similar pockets, based on the observation that structure-similar pockets often exhibit similar interaction patterns with ligands. The aggregation function is defined as:

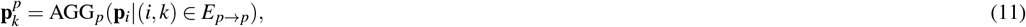

where 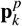 is the refined embedding of the *k*-th pocket **p**_*k*_ that aggregates information from its neighboring pockets; AGG_*p*_ is implemented using an attention mechanism, which is defined as follows:

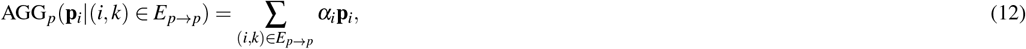

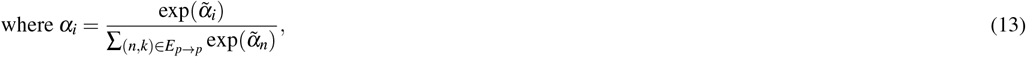

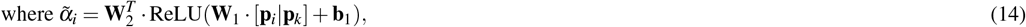

where *α*_*i*_ denotes the normalized attention weight for the neighboring pocket *i*; [**p**_*i*_ | **p**_*k*_] denotes the concatenation of vectors **p**_*i*_ and 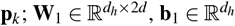, and 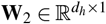 are three learnable parameters, where *d* is the hidden dimension of pocket embeddings and *d*_*h*_ is the hidden dimension of the attention network.

### Pocket-ligand aggregator

After aggregating information from similar pockets, We next utilize the pocket-ligand aggregator to aggregate information from 2-hop ligand neighbors, based on the observation that structure-similar pockets should have structure-similar active ligands. Let 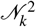 denote the set of 2-hop ligand neighbors of pocket *k*:

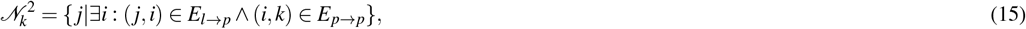

the aggregation function is defined as:

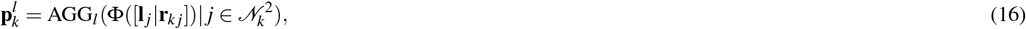

where 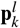 is the refined embedding of the *k*-th pocket that aggregates information from the active ligands of its neighbor pockets; **l** _*j*_ is the ligand embedding of the *j*-th ligand; **r**_*kj*_ is a relation embedding derived from the similarity score between the *k*-th pocket and its intermediate neighbor (the *i*-th pocket) where the *j*-th ligand is active for it; Φ is a 2-layer MLP with the hidden size of 2*d*. Similar to the pocket-pocket aggregator, AGG_*l*_ is also implemented using an attention mechanism with different learnable parameters.

### Refined pocket embedding

Finally, we combine the original pocket embedding with the refined embeddings from both aggregators to obtain the final refined pocket embedding:

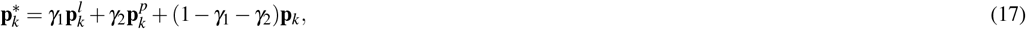

where *γ*_1_, *γ*_2_ *∈* [0, 1] and *γ*_1_ + *γ*_2_ *≤* 1 are parameters that control the contribution of each component; this weighted combination allows the model to adaptively balance information from different sources: similar pockets, their active ligands, and the pocket’s intrinsic features. Collectively, our proposed H-GNN integrates information from both similar pockets and their active ligands, leveraging the observation that structure-similar binding pockets tend to have similar active ligands, to enhance the performance of LigUnity in finding active ligands for the query protein pocket. The effectiveness of H-GNN is shown in our ablation studies (**Supplementary Figures. 11-13**).

## Resource availability

The datasets for training LigUnity were collected from ChEMBL version 34^21^ and BindingDB version 2024m5^20^. Our training dataset is available on figshare https://doi.org/10.6084/m9.figshare.27966819. Our PocketAffDB with protein and pocket PDB structures is available on figshare https://doi.org/10.6084/m9.figshare.29379161. Our processed Dekois 2.0 benchmark dataset is available on figshare https://doi.org/10.6084/m9.figshare.27967422. The LIT-PCBA and DUD-E benchmark datasets were obtained from Gao et al. https://github.com/bowen-gao/DrugCLIP^13^. The FEP calculation benchmarks^9,31^ and the calculated results using FEP+(OPLS4)^9^ were obtained from https://github.com/schrodinger/public_binding_free_energy_benchmark^63^. The experimental affinity data for the PDE10A data released by Roche Ltd. was obtained from Tosstorff et al.^35^ The binding free energy dataset on the TYK2 target released by Google Research was obtained from Thompson et al.^36^

The code is available in our GitHub project https://github.com/IDEA-XL/LigUnity, including the model weights of LigUnity and relevant source code. We have included details of our experiments in the Methods to enable independent replication.

## Author contributions

B.F., Z.L., M.Y., Y.L., L.Z., and S.W. contributed to the conception and design of the work. B.F. and Z.L. contributed to the data curation. B.F. and Y.L. contributed to the technical implementation. B.F., Z.L., H.L., Y.L., L.Z., and S.W. contributed to technical discussions. B.F., Z.L., M.Y., J.Z., and S.W. contributed to the evaluation framework used in the study. All authors contributed to the drafting and revision of the manuscript.

## Acknowledgments

We’d like to express our gratitude to Dr. Zequn Liu, Dr. Kangjie Zheng, and Dr. Junwei Yang from Peking University for their valuable discussions and assistance in this work. We are also grateful to Dr. Michael K. Gilson and Dr. Tiqing Liu from the University of California, San Diego (UCSD) for their expert guidance on data curation from BindingDB.

This project was supported by Shenzhen Hetao Shenzhen-Hong Kong Science and Technology Innovation Cooperation Zone, under Grant No. HTHZQSWS-KCCYB-2023052.

## Supplemental information

Document S1. Implementation details, Data curation, Competing methods, Experimental settings, Supplementary Figure 1-21, and Supplementary Tables 1-4.

## Supplementary information

### Implementation details

The details of our training strategy are listed as follows. In the pre-training stage, we trained LigUnity for 50 epochs, and we used the averaged parameter of the last 10 epochs for testing. We warmed up the learning rate for the first 6% step and adopted a polynomial decay learning rate scheduler afterward. We used the Adam optimizer with a maximum learning rate of 0.0001. For each training iteration, we sampled 24 assays for computing loss. For each assay, we randomly sampled one pocket structure and a maximum of 16 measured ligands coming from the assay, making sure that their experimental affinity values were comparable to each other. We used listwise ranking for measured ligands that belonged to the same assay and used contrastive loss between the whole sampled ligands and pockets in this batch.

For Heterogeneous-GNN (H-GNN), we trained it separately from the pocket and ligand encoder. We trained H-GNN for 10 epochs and used the last epoch for testing. We adopted the constant learning rate scheduler with a learning rate of 0.001. For each training iteration, we randomly sampled 128 assays, and we randomly selected one pocket structure and one active ligand for each assay. As the large-scale nature of the pocket-ligand knowledge graph makes it hard to fine-tune H-GNN on a few measured ligands, we adopted H-GNN only for virtual screening tasks and used the contrastive loss between the whole ligands and pockets sampled in this batch for training.

For LigUnity(seq), the protein-based version of LigUnity, we used ESM2^64^ 35M for encoding the protein sequence in replacement of the 3D pocket encoder, and we updated the parameters of ESM2 during training. Furthermore, we also attempted to use the larger ESM2 3B model with a frozen gradient but observed a drop in test performance. To fully make use of the pocket and protein information, the results of LigUnity are reported as the ensemble of the pocket-based model and the protein-based model, except for the experimental setting where training proteins with >30% sequence similarity are removed. This exception is made because protein-based models typically show limited generalization ability to unseen protein families with low sequence similarity.

We also trained Pocket-DTA and Sequence-DTA, the regression-based version of LigUnity and LigUnity(seq) respectively, to compare different protein-ligand affinity prediction strategies. In the pre-training stage, we used mean square loss and margin loss for training. First, for pocket-ligand pairs with experimental affinities, we used the mean square loss for training which is defined as the mean square error between experimental affinity and predicted affinity (*L*_*SE*_ = (*y*_*i*_− *ŷi*)^2^) . Second, to enable Pocket-DTA to distinguish between active and inactive ligands, we introduced in-batch margin loss. For unmeasured in-batch pocket-ligand pairs, we consider them to be inactive and employ margin loss to penalize them (*L*_*Margin*_ = max(0, *ŷ* − (*y*_*min*_ − *m*))^2^), where *y*_*min*_ is the minimum known activity known to the pocket and *m* is a margin hyper-parameter. We set the margin to 2.0 after a grid search on the validation set.

### Data curation

The purpose of our data curation is to collect high-quality pocket-ligand binding affinity data and organize them by assays. We developed a curation pipeline based on the BindingDB^20^ and ChEMBL^21^ datasets. ChEMBL is a large-scale bioactivity database that integrates experimental data from scientific publications, patents, and other datasets. It covers various aspects of drug discovery and development, containing 2.5 million unique compounds and 19 million experimentally measured activities. BindingDB is another large-scale database that focuses specifically on protein-ligand binding affinities. It is curated mainly from scientific publications and US patents, containing 0.5 million compounds, 1.1 million measured affinities, and 2,773 protein targets.

Our data curation pipeline consists of three stages: organizing affinities into assays, assay-guided pocket matching, and removing duplicate assays. In this study, we used the 34 version of ChEMBL and the 2024m5 version of BindingDB downloaded from their official websites. The details of our curation pipeline are as follows:

### Organizing affinities into assays

Considering that the affinities from different assays are incompatible due to varying experimental conditions (e.g., cofactor concentration, pH, temperature), assay formats (e.g., cell-based vs. target-based assays, and different detection methods), and affinity measurements (e.g., IC50, Ki, Kd), we first organized the experimental affinities into assays. For ChEMBL, we collected all assays with “Binding” type and known protein targets, then we grouped affinity data by ChEMBL assay ID. We only retained affinity data with molar concentration units (e.g., nmol) or density units (e.g., *µ*g*/*ml) to ensure data quality, as the exact meaning of other units (e.g., % or no unit) was difficult to determine. This filtering step removed 33.1% of the activity data in ChEMBL. For BindingDB, we grouped the affinity data with the same protein target, affinity measurement, source document, and assay description into an assay following Actfound^14^, ensuring that affinity values within the same assay were comparable. We also removed inorganic ligands and large molecular ligands (molecular weights *>* 1,000). Finally, we retained assays with more than 5 affinity values for further curation. We collected 69,872 and 25,902 assays from ChEMBL and BindingDB respectively, with each assay having one corresponding protein target.

### Assay-guided pocket matching

Based on the observation that most assays are designed for a specific binding site of interest, we hypothesize that ligands in an assay should bind to the same pocket of a protein. Therefore, once we identified a binding pocket for one ligand in an assay, we assumed that other ligands in the same assay would bind to the identical pocket. For each assay, we identified binding pockets in the PDB (Protein Data Bank)^65^ using two criteria: protein sequence similarity and ligand Tanimoto similarity. First, we searched for protein-ligand complex structures whose protein sequence similarity with the protein studied in this assay is above 40%, as proteins with such similarity typically share high structural and functional homology^33^. Then, we calculated the Tanimoto similarity between the crystal ligands in these PDB structures and the assay ligands. If the ECFP4 Tanimoto similarity exceeded 70%, we collected the binding pocket of the crystal ligand as the candidate pockets for this assay, as ligands with such high similarity are likely to bind to the same pocket in a protein^66^. We have trained model on dataset curated with 100% Tanimoto similarity threshold and observed a performance decrease (**Supplementary Table. 4**).

To discard biologically irrelevant pockets that have weak binding interactions with the ligand (i.e., those having a small number of binding residues with ligands), we used the processed binding pocket database provided by BioLip^67^. After this procedure, we collected 25,883 and 16,863 assays from ChEMBL and BindingDB respectively. For each assay, there are 7.19 identified pockets on average. Although the identified pockets for the same assay have different PDB IDs, they mostly belong to the same binding site. To verify this, we aligned complex PDB structures retrieved for each assay and computed the maximum distance between the centers of all pockets. We identified 1,229 and 802 assays whose maximum distance between retrieved pockets exceeded 10 Å for ChEMBL and BindingDB respectively, accounting for only 4.8% of the total assays. We have trained model on assays without multiple pockets (>10Å) and observed a performance decrease (**Supplementary Table. 4**).

### Removing duplicate assays

As ChEMBL and BindingDB share common data sources including scientific literature and patents, there could be many overlapping assays between these two datasets. Additionally, there are some repeating assays within the ChEMBL or BindingDB database as well. To reduce the negative impact of data redundancy on pre-training, we detected and removed identical assays following pQSAR-ChEMBL^68^. Specifically, if the Pearson correlation (*ρ*_*p*_) of affinities between the shared ligands of two assays exceeds a threshold of 0.95, we considered those two assays to be identical. After this stage, we detected and removed 15,664 repeating assays, ending up with 26,748 non-repeating assays curated from ChEMBL and BindingDB.

After the above curation procedures, we collected 26,748 assays with 428,767 unique ligands and 36,662 unique pockets from ChEMBL, BindingDB, and PDB, covering 2,196 unique proteins. Each assay has on average 30.29 measured ligands and 10.96 pockets from PDB. Our curation process has created a large-scale pocket-ligand affinity dataset while maintaining the quality of the curated protein binding pockets, which is supported by the small fraction of assays with distinct pockets.

However, due to the limited number of unique proteins in the original binding database (2,773 in BindingDB), models trained solely using these data might result in poor generalization on unseen proteins. To address this limitation, we further enlarged our curated dataset by utilizing the protein-ligand complex data from PDB^69^, which includes diverse proteins. For each protein-ligand complex with crystal structure, we considered the ligand to be active following previous methods^13,27^ and created one assay that contained only one protein pocket and one active ligand. We did not merge ligands binding to the same protein into one assay, as we could not determine the relative affinity ranking between these ligands. To avoid the negative impact of the low-quality complex structure data in PDB (those with low resolution, inactive ligands, or biologically irrelevant ligands), we directly used the PDBBind v2020^70^ dataset for training, following previous methods^13,27^. The PDBBind dataset contains 3,889 unique proteins, and the number of unique proteins and pockets in our curated pre-training data finally reached 4,847 and 53,406 respectively.

### Competing methods

In this paper, we compared LigUnity with three types of competing methods: (1). docking score methods including Glide (SP and XP)^25^, Vina^26^, AutoDock4^71^, Vinardo^72^, Gold^73^, Surflex^74^, and Flexx^75^; (2). structure-based ML methods that take protein-ligand complex structure as input including NNscore 2.0^48^, RFscore v4^49^, Δ_Vina_RF^50^, Δ_Lin_F9_XGB^52^, RTMScore^27^, Genscore^16^, Pafnucy^53^, OnionNet^54^, Planet^56^, PIGNet^55^, EquiScore^19^, Kdeep^76^, TANKBind^77^, 3D-GNN^55^, PBCNet^15^, Denvis^28^ and GNINA^57^; (3). structure-free methods DeepDTA^29^, BIND^78^, DrugCLIP^13^, Pocket-DTA (the regression-based version of LigUnity) and its protein sequence-based version Sequence-DTA. For experiments on the Merck and JACS benchmarks, we additionally compared LigUnity with physics-based binding free energy calculation methods including Prime-MM/GBSA^32^ and FEP+^9^. For active learning experiments on the TYK2 dataset, we compared five machine learning methods including ElasticNet^79^, Gaussian Process^80,81^, Gradient Boosting Machine (GBM)^82^, Linear Regression, and Random Forest^83^, and these methods take 2048-dimensional ECFP4 fingerprint as input.

### Experiments on virtual screening benchmarks

For virtual screening, we evaluated our method on three benchmark datasets (DUD-E^22^, Dekois^23^, and LIT-PCBA^24^), all of which have confirmed binding sites. For DUD-E and LIT-PCBA benchmarks, we used the processed version from DrugCLIP’s repository^13^. For the Dekois 2.0 benchmark, we downloaded it from the official website and extracted residues within 6 Å distance from the crystal ligand to form the binding pocket. To prevent data leakage from overlapping protein targets between our curated pre-training data and test benchmarks, we excluded proteins that appear in the three benchmarks from our pre-training data and used the model trained on this processed dataset for testing.

The DUD-E benchmark contains 102 targets, with an average of 224.4 active ligands per target. For each active ligand, there are approximately 50 decoys with similar physicochemical properties, with over 99% of decoys not experimentally verified. The Dekois 2.0 benchmark follows a similar construction approach, containing 81 targets, each with 30 active ligands and 40 generated decoys per active ligand. The LIT-PCBA benchmark has 15 targets with an average of 533.3 active ligands per target. It contains 2.64 million inactive ligands with an active-to-inactive ratio of 1:1000, better reflecting real-world drug design scenarios. Additionally, all active and inactive ligands in LIT-PCBA have been experimentally verified. In contrast, DUD-E and Dekois 2.0 benchmarks exclude decoys with similar structures to avoid potential active ligands, which may make them relatively easy for ML models. For DUD-E, the results of Denvis-G, Denvis-R, DeepDTA, Gold, Surflex, Flexx, Vina, and GNINA are from Krasoulis et al.^28^; the results of other structure-based methods are from Cao et al.^19^. For Dekois 2.0, the results of all docking methods and structure-based methods are from Cao et al.^19^. For LIT-PCBA, the evaluation results of Denvis-G, Denvis-R, GNINA, and DeepDTA are from Krasoulis et al.^28^; the results of GenScore and Glide SP are from Shen et al.^16^

Following previous works^13,27,28^, we evaluated methods using three metrics: Enrichment Factor (EF), Area Under the Receiver Operating Characteristic curve (AU-ROC), and Boltzmann-Enhanced Discrimination of ROC (BEDROC). EF quantifies a model’s ability to identify actives among top-ranked ligands, calculated as the proportion of actives in top-ranked ligands (e.g., top 1%) divided by the proportion of actives in all candidate ligands. We use EF 1% for evaluation, which ranges from 0 to 100. AU-ROC assesses a model’s ability to correctly rank active ligands before inactive ligands and is defined as the area under the plot of true positive rate against false positive rate. However, since AU-ROC treats all ranked ligands equally, it may not be ideal for virtual screening tasks where only top-ranked ligands undergo costly experimental verification. To address this limitation, we employ BEDROC, which assigns higher weights to top-ranked ligands. Following previous works^13,27,28^, we set the *α* parameter to 80.5 to balance the evaluation of top-ranked ligands and overall ranking performance. AU-ROC and BEDROC both range from 0 to 1. For all metrics, higher values indicate better performance.

To further assess LigUnity’s generalizability to unseen proteins, we conducted experiments on unseen protein families. We excluded training proteins with more than 80% and 30% sequence similarities to any test protein on the three benchmarks; then we used the model trained on this processed dataset for evaluation. We used cd-hit to compute protein sequence similarity, which is defined as the number of identical residues after alignment.

### Experiments on FEP benchmarks

To test the performance of LigUnity on hit-to-lead optimization scenarios, we evaluated all methods on two FEP calculation benchmarks (JACS^9^ and Merck^31^), all of which have confirmed binding sites. The JACS and Merck benchmark datasets, along with the results of FEP+(OPLS4)^9^ calculated using the state-of-the-art OPLS4^84^ force field, were obtained from the public repository https://github.com/schrodinger/public_binding_free_energy_benchmark^63^. These FEP benchmarks contain 16 targets (assays), with each target having an average of 29 experimentally measured ligands. We extracted residues within a 6 Å distance from the crystal ligand as the binding pocket. For experiments on these FEP calculation benchmarks, we used Spearman correlation (*ρ*_*s*_) and *r*^2^ = max(0, *ρ*_*p*_)^2^ for evaluation.

For zero-shot evaluation on FEP benchmarks, we directly applied LigUnity for inference without fine-tuning. To mitigate potential data leakage in the pre-training data, we removed assays that were similar to assays in test benchmarks from the pre-training data. Following previous methods^14,68^, we defined two assays as identical if the affinities of overlapping ligands reached a Pearson’s correlation of 0.95. We also removed all ligands from the pre-training data that existed in FEP benchmarks to ensure a zero-shot setting. To reduce variability in evaluation metrics due to the limited number of targets in the FEP benchmarks, we trained five models with different random seeds for all our implemented methods (LigUnity, LigUnity(seq), Pocket-DTA, and Sequence-DTA), and reported the average evaluation metrics. The evaluation results (*ρ*_*s*_) of all computational methods and structure-based methods on the Merck benchmark are from Shen et al.^16^

For few-shot experiments, the fine-tuning ligands for each test assay were randomly sampled with a uniform distribution, and we evaluated LigUnity’s performance on the remaining ligands. We evaluated all methods with varying proportions of fine-tuning ligands ranging from 20% to 80%. Following previous methods^14^, we repeated the test experiments 40 times. For all our implemented methods (LigUnity, LigUnity(seq), Pocket-DTA, and Sequence-DTA) and DrugCLIP, we performed 5 steps of fine-tuning on the fine-tuning ligands using the same learning rate as in pre-training, and the final evaluation metrics were obtained by averaging the model’s predictions on test ligands from all steps.

To study the generalization ability of LigUnity on unseen chemical scaffolds and proteins, we investigated three increasingly challenging settings: (1) no similar ligands setting, where we excluded training ligands with more than 50% ECFP4 Tanimoto similarity to any test ligand, the similarity is computed using 2048-dimensional ECFP4 fingerprints generated by RDKit; (2) no similar proteins setting, where we excluded training proteins with more than 40% sequence similarity to any test protein (threshold set following AlphaFold3^33^), and we used cd-hit to compute protein sequence similarity defined as the number of identical residues after alignment; (3) no similar ligands and proteins setting, where we applied both criteria from the first two settings.

For calculating the importance score of each ligand atom and protein residue, we replaced atomic features with padding tokens and measured the resulting decrease in affinity prediction, where larger decreases indicated higher importance of the masked atoms. In this study, we used the model trained on the “no same ligands” setting, and we used the PDB structure 5EHR and its crystal ligand 5OD for the case study.

### Experiments under diverse settings

After evaluation on FEP benchmarks, we next sought to evaluate LigUnity on assays from ChEMBL^21^ and BindingDB^20^, which contain diverse assays covering various aspects of assay types. We evaluated all methods under two assay split settings: split-by-time and split-by-unit. Under each assay split setting, we also investigated two ligand split settings: uniformly split at random and split-by-scaffold.

For the split-by-time setting, we first pre-trained all models using assays released before March 2019, and then conducted testing on assays released after March 2019. To evaluate generalization capability on truly unseen proteins, we only evaluated on assays containing previously unseen proteins. This test set comprised 161 assays, with an average of 48.6 ligands per assay. For the split-by-unit setting, we collected 65 assays with unit “%” and assay type “Activity” from ChEMBL for testing, differing from our pre-training assays that have molar concentration units (e.g., nmol) or density units (e.g., *µ*g*/*ml). This test set contains 65 assays in total, with an average of 30.5 ligands per assay. In this setting, we used the model trained in the split-by-time setting for evaluation.

For few-shot evaluation on both split-by-time and split-by-unit settings, we evaluated all methods with varying proportions of fine-tuning ligands ranging from 4 to 16. We repeated the test experiments 10 times. For all our implemented methods (LigUnity, LigUnity(seq), Pocket-DTA, and Sequence-DTA) and DrugCLIP, we performed 10 steps of fine-tuning on the fine-tuning ligands using the same learning rate as in pre-training, and the final evaluation metrics were obtained by averaging the model’s predictions on test ligands from the last 5 steps (steps 6-10). For the split-by-scaffold setting on each test assay, we first generated Bemis-Murcko chemical scaffolds for all ligands using RDKit; then we iteratively selected molecules sharing the same scaffold as fine-tuning ligands until reaching the preset number, while using the remaining molecules for testing.

For evaluation on the PDE10A target, we used the dataset released by Roche Ltd. in 2022^35^. This dataset comprises 1,162 ligands with experimentally measured binding affinities and corresponding experimental timestamps. To simulate the realistic drug development scenarios, we followed the split-by-time setting at both assay and ligand levels (**Fig. 4e**). At the assay splitting level, we used the model pre-trained on assays published before March 2019, while the PDE10A dataset was released in 2022. At the ligand splitting level, we used 20% of the ligands measured earlier for fine-tuning and the remaining 80% of ligands measured later for testing. As the number of fine-tuning ligands is more than 100, we performed 20 steps of fine-tuning, and the final evaluation metrics were obtained by averaging the model’s predictions on test ligands from the last 5 steps (steps 16-20).

### Active learning experiment on the TYK2 target

For the active learning experiment on the TYK2 target, we used the TYK2 kinase inhibitor dataset released by Google Research^36^. This dataset contains 10,000 ligands with binding free energies calculated using FEP. The dataset was initially constructed from 573 measured ligands found in public patents and was then expanded to 10,000 ligands through R-group assembly and filtering. The binding free energies were calculated using the Relatively Absolute Binding Free Energy (RABFE) protocol^85^, which achieved an *R*^2^ of 0.41 with experimental affinities in validation tests. The entire calculation consumed 80,000 GPU hours (equivalent to 9.1 GPU years), which would cost 160,000$.

We implemented an active learning framework to evaluate LigUnity and competing methods. Our goal was to identify ligands with optimal binding free energy within specific FEP calculation constraints. The procedure consisted of the following steps: (1) Sampling a pool of ligands using a defined selection strategy; (2) Calculating binding free energies for selected ligands using the FEP method and incorporating them into the training set (in this study, we directly used existing values from the dataset); (3) Training a model using ligands with known binding free energies; (4) Using the trained model to predict binding affinities for unmeasured ligands; (5) Repeating steps 1-4 until reaching a predetermined number of accessed ligands. This procedure simulates real-world drug discovery scenarios, providing insights into the practical utility of different methods.

For each iteration, we explored 100 ligands, starting with a randomly sampled initial training set of 100 ligands, and we conducted 20 active learning iterations in total. For each iteration, we performed 20 fine-tuning steps on the fine-tuning ligands. The final predictions were obtained by averaging the model’s predictions on test ligands from the last 5 steps (steps 16-20). We evaluated each method using two types of metrics: (1) top 1%, 2%, and 5% recall, defined as the ratio of correctly identified top ligands to the total number of top ligands, and (2) *r*^2^, calculated based on the ligands in the test set.

**Supplementary Figure 1:**
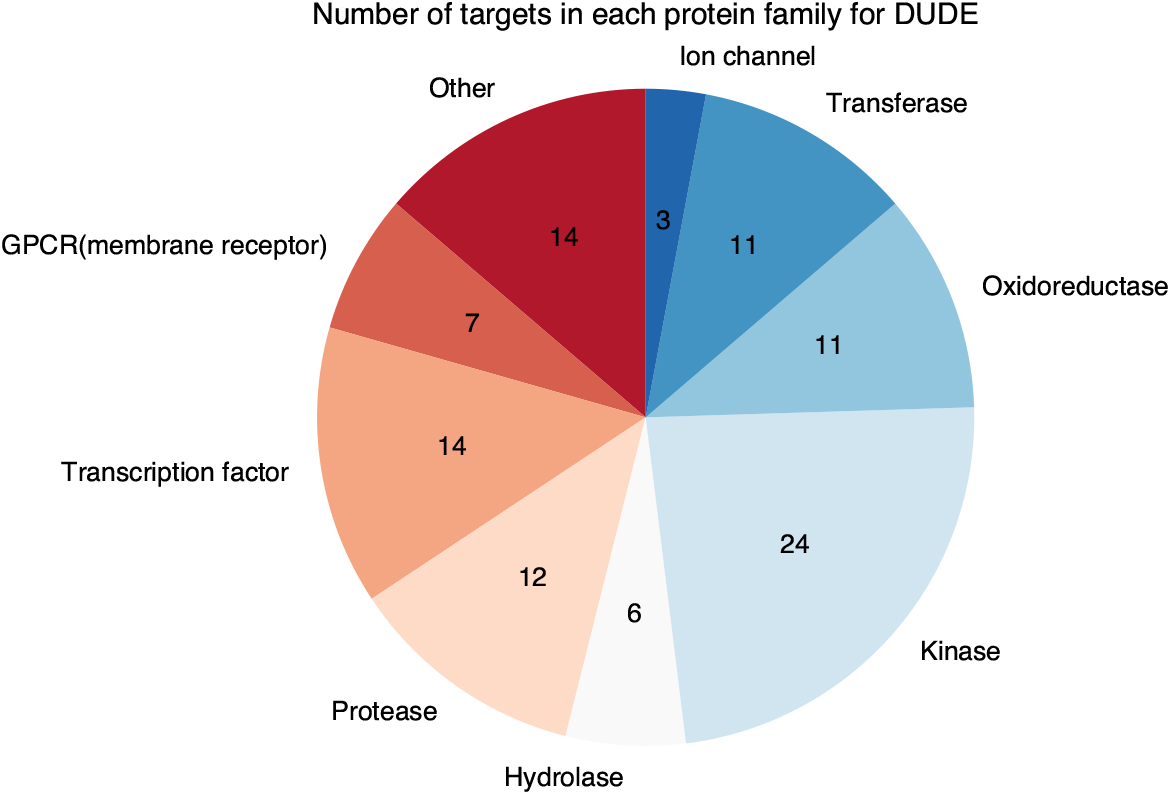
Pie chart showing the number of targets for each protein family in the DUD-E benchmark. The protein families come from ChEMBL^21^.

**Supplementary Figure 2:**
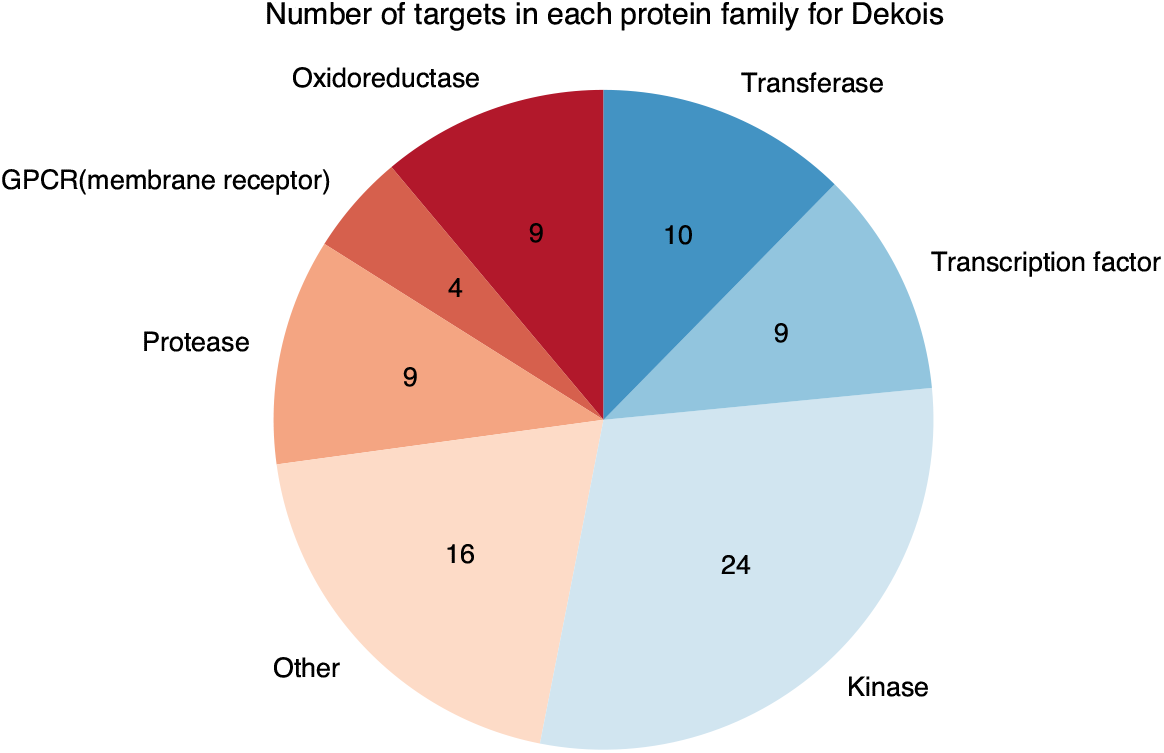
Pie chart showing the number of targets for each protein family in the Dekois 2.0 benchmark. The protein families come from ChEMBL^21^.

**Supplementary Figure 3:**
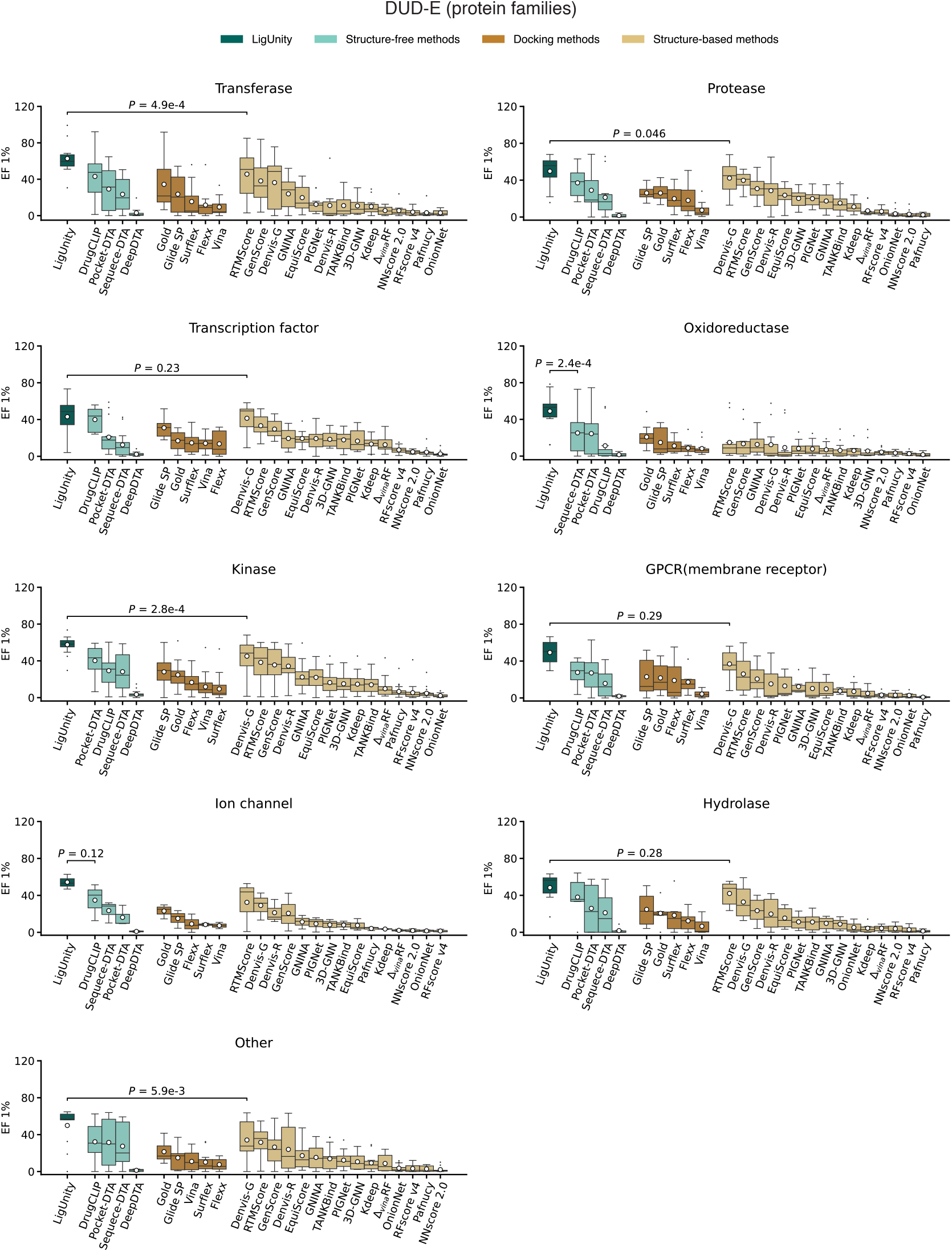
Box plots comparing LigUnity and competing methods on the DUD-E benchmark in terms of enrichment factor (EF) 1%. Targets of different protein families are tested separately.

**Supplementary Figure 4:**
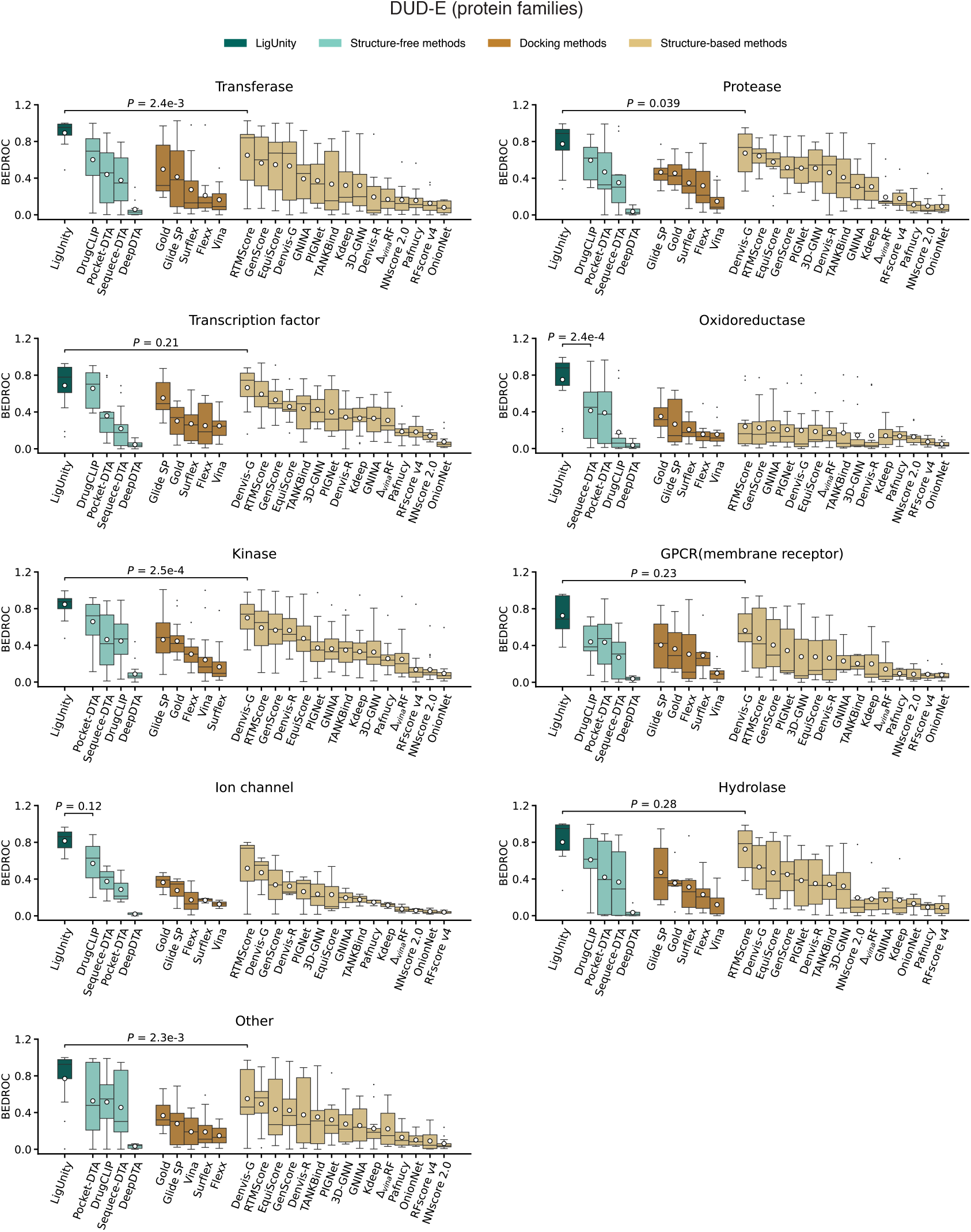
Box plots comparing LigUnity and competing methods on the DUD-E benchmark in terms of BEDROC. Targets of different protein families are tested separately.

**Supplementary Figure 5:**
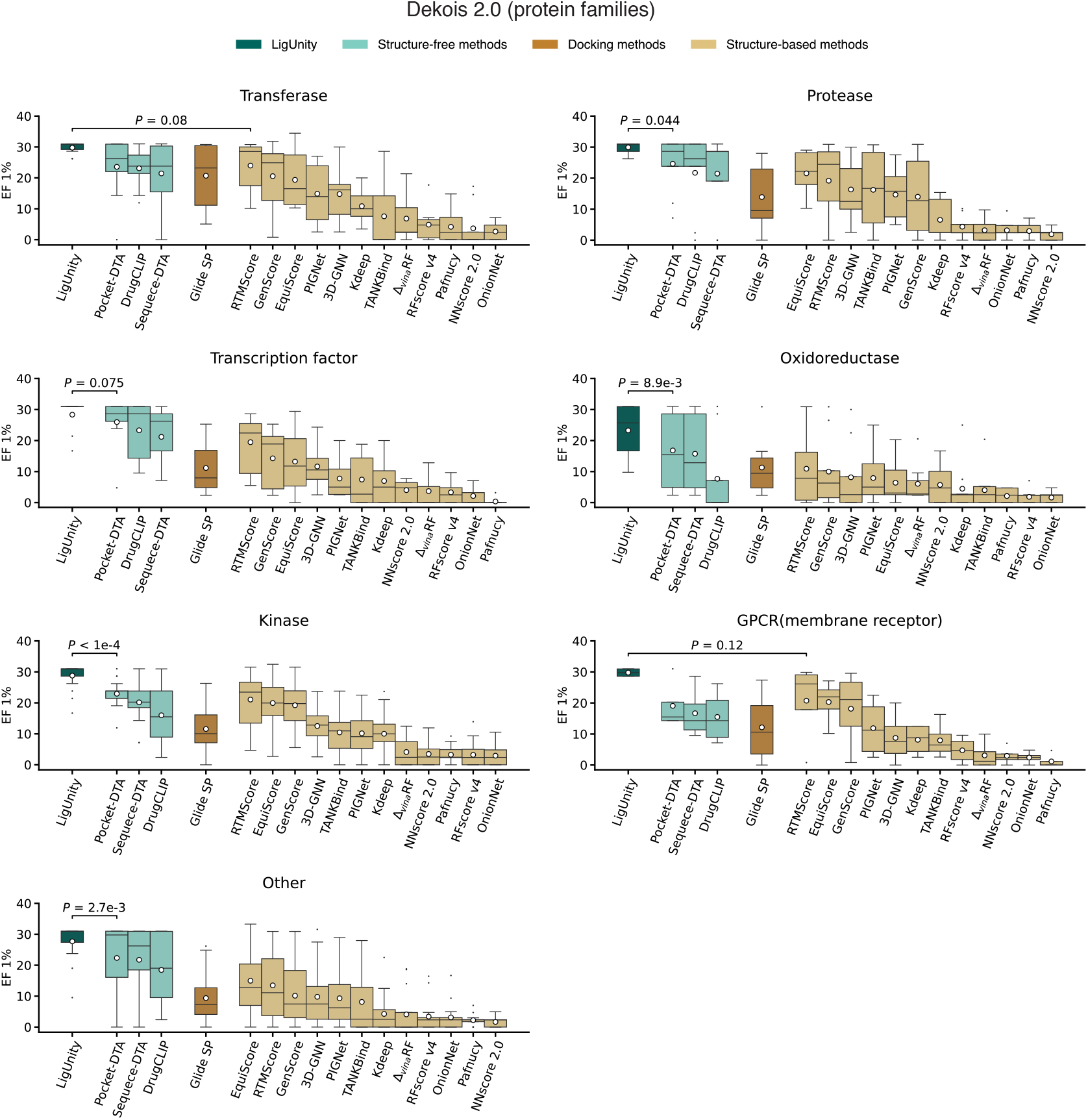
Box plots comparing LigUnity and competing methods on the Dekois 2.0 benchmark in terms of enrichment factor (EF) 1%. Targets of different protein families are tested separately.

**Supplementary Figure 6:**
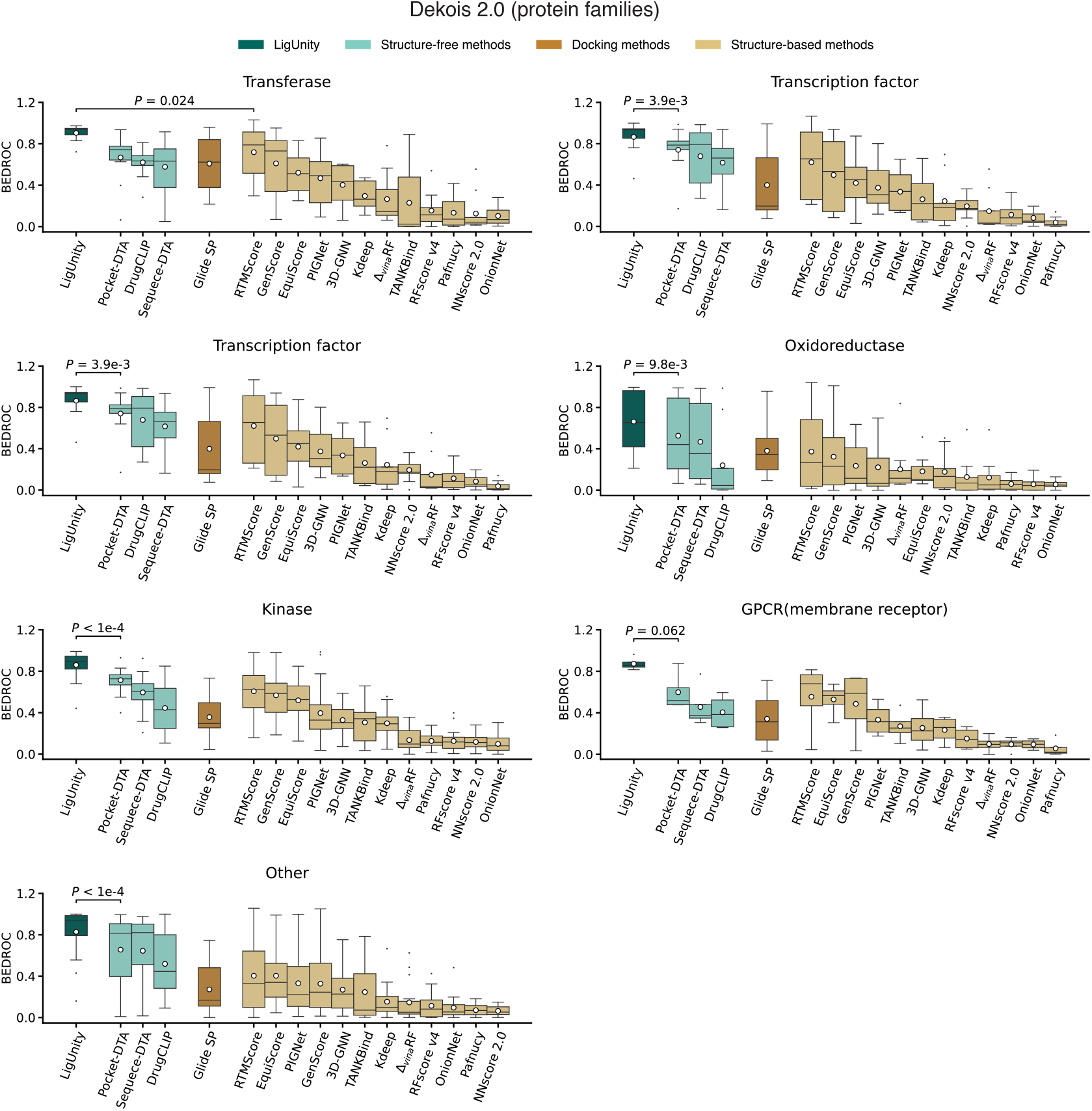
Box plots comparing LigUnity and competing methods on the Dekois 2.0 benchmark in terms of BEDROC. Targets of different protein families are tested separately.

**Supplementary Figure 7:**
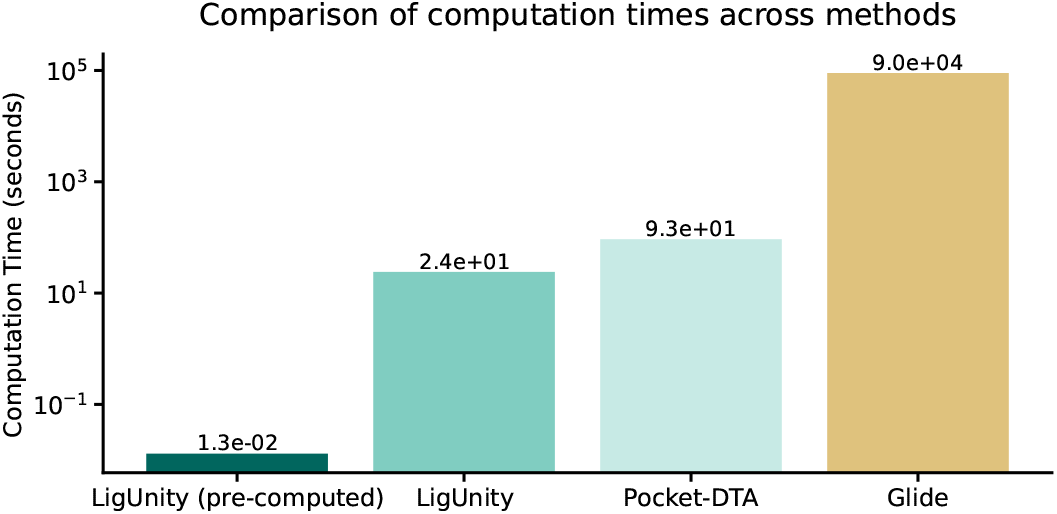
Bar plots compare LigUnity and competing methods based on the computation times for screening 10,000 ligands. Glide-SP’s computation time is estimated on a 2.8 GHz Intel Xeon E5-1603 processor, while LigUnity’s and Pocket-DTA’s times are estimated on a NVIDIA RTX A6000 GPU.

**Supplementary Figure 8:**
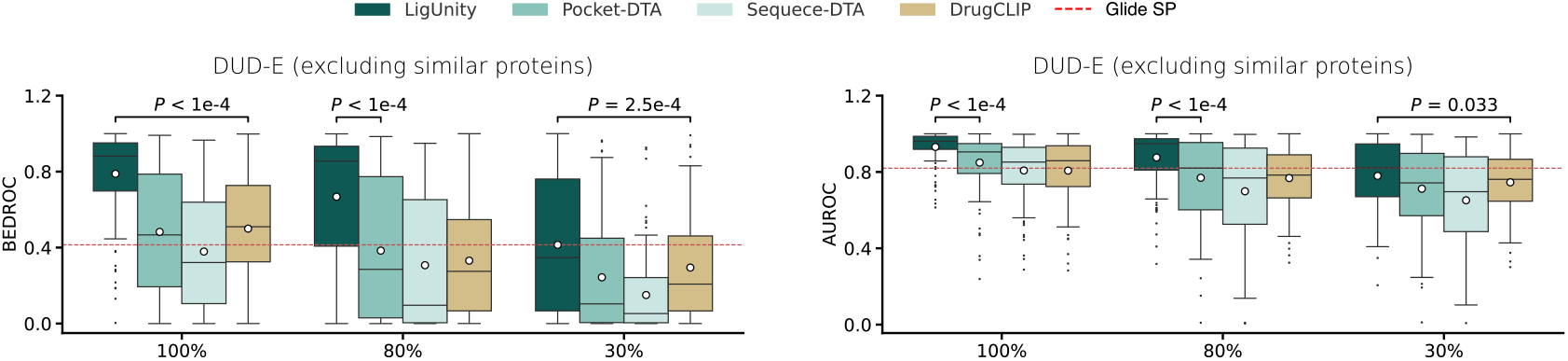
Box plots comparing LigUnity and competing methods on the DUD-E benchmark in terms of BEDROC and AU-ROC using different protein training sets. The *x*-axis denotes the maximum sequence similarity between the training and test sets. The mean values (white dots) are calculated across n = 102 targets.

**Supplementary Figure 9:**
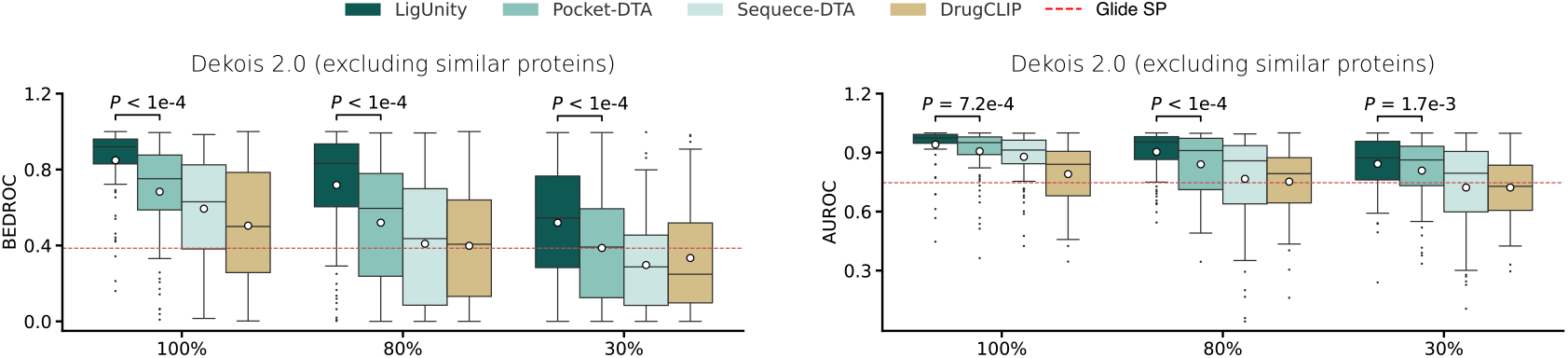
Box plots comparing LigUnity and competing methods on the Dekois 2.0 benchmark in terms of BEDROC and AU-ROC using different protein training sets. The *x*-axis denotes the maximum sequence similarity between the training and test sets. The mean values (white dots) are calculated across n = 81 targets.

**Supplementary Figure 10:**
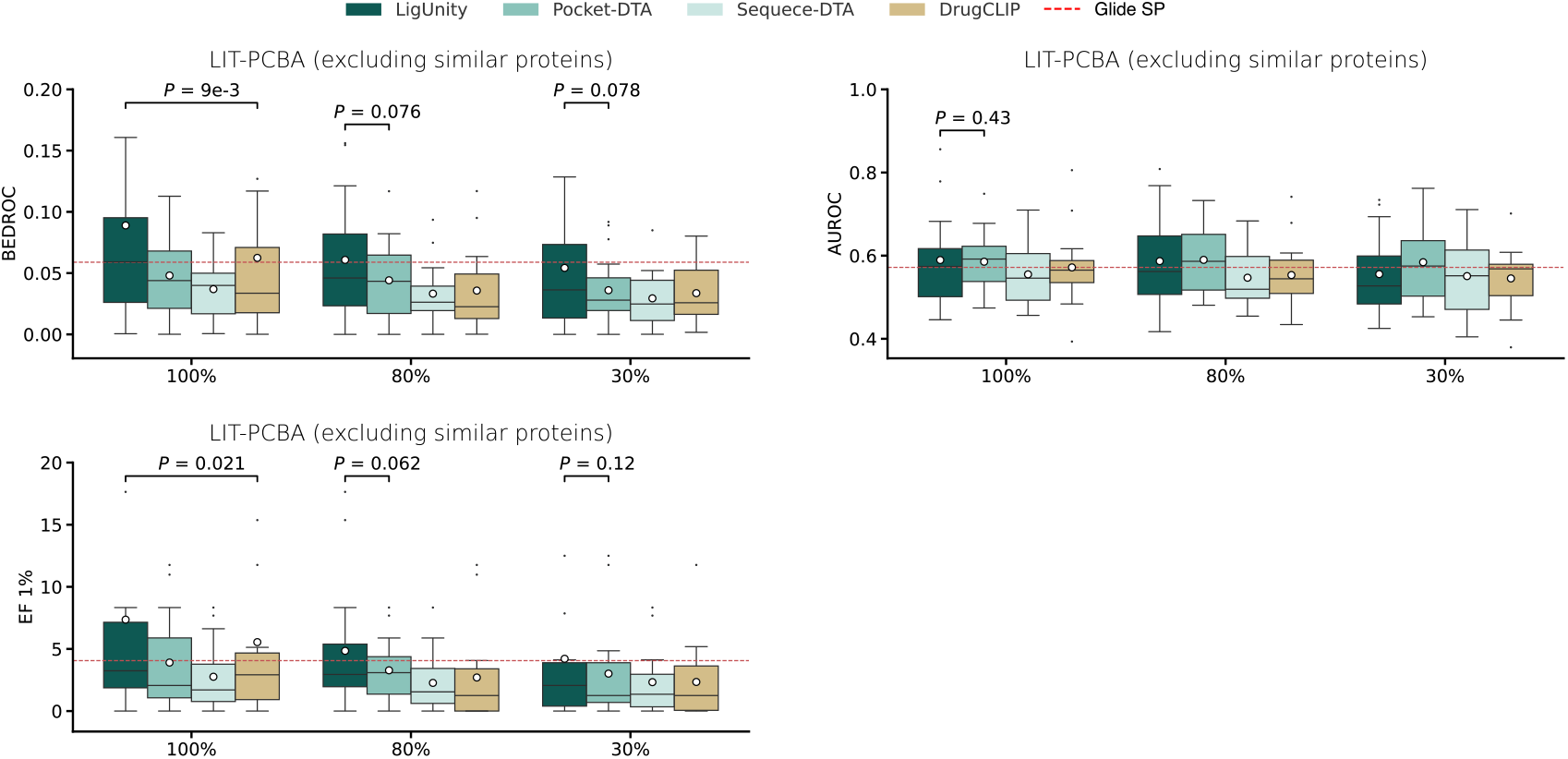
Box plots comparing LigUnity and competing methods on the LIT-PCBA benchmark in terms of enrichment factor (EF) 1%, BEDROC, and AU-ROC using different protein training sets. The *x*-axis denotes the maximum sequence similarity between the training and test sets. The mean values (white dots) are calculated across n = 15 targets.

**Supplementary Figure 11:**
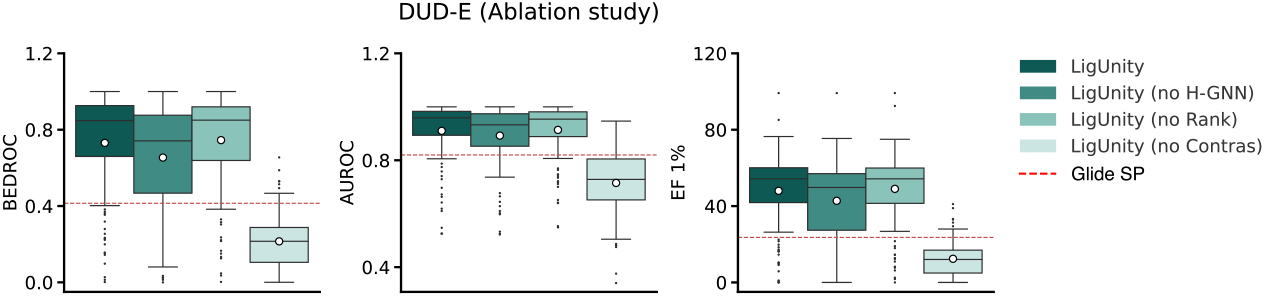
Box plots of ablation study evaluating the contribution of different modules on the DUD-E benchmark in terms of enrichment factor (EF) 1%, BEDROC, and AU-ROC, where “no Contras” means that the contrastive loss for scaffold discrimination are ablated and “no Rank” means that the ranking loss for pharmacophore ranking are ablated. The mean values (white dots) are calculated across n = 102 targets.

**Supplementary Figure 12:**
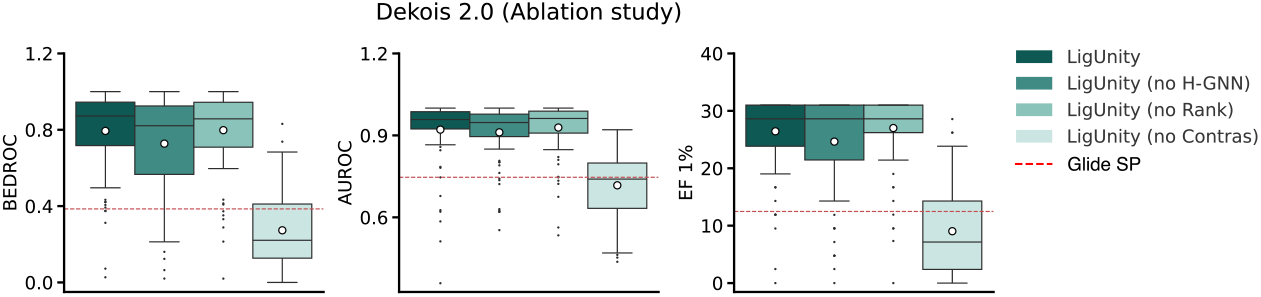
Box plots of ablation study evaluating the contribution of different modules on the Dekois 2.0 benchmark in terms of enrichment factor (EF) 1%, BEDROC, and AU-ROC, where “no Contras” means that the contrastive loss for scaffold discrimination are ablated and “no Rank” means that the ranking loss for pharmacophore ranking are ablated. The mean values (white dots) are calculated across n = 81 targets.

**Supplementary Figure 13:**
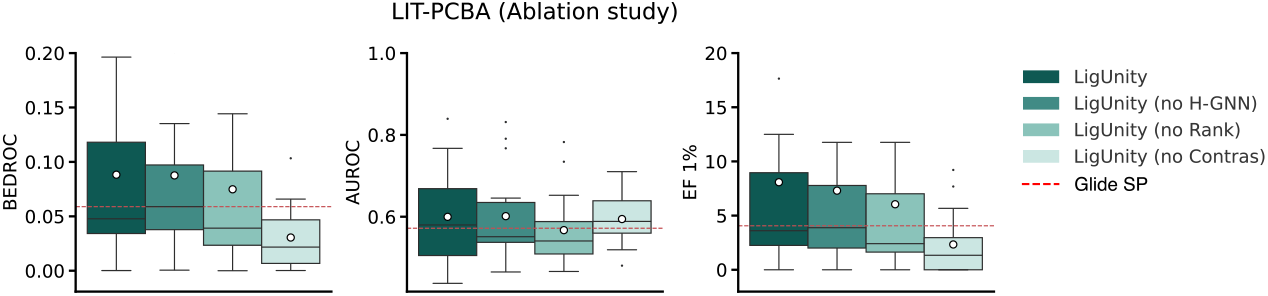
Box plots of ablation study evaluating the contribution of different modules on the LIT-PCBA benchmark in terms of enrichment factor (EF) 1%, BEDROC, and AU-ROC, where “no Contras” means that the contrastive loss for scaffold discrimination are ablated and “no Rank” means that the ranking loss for pharmacophore ranking are ablated. The mean values (white dots) are calculated across n = 15 targets.

**Supplementary Figure 14:**
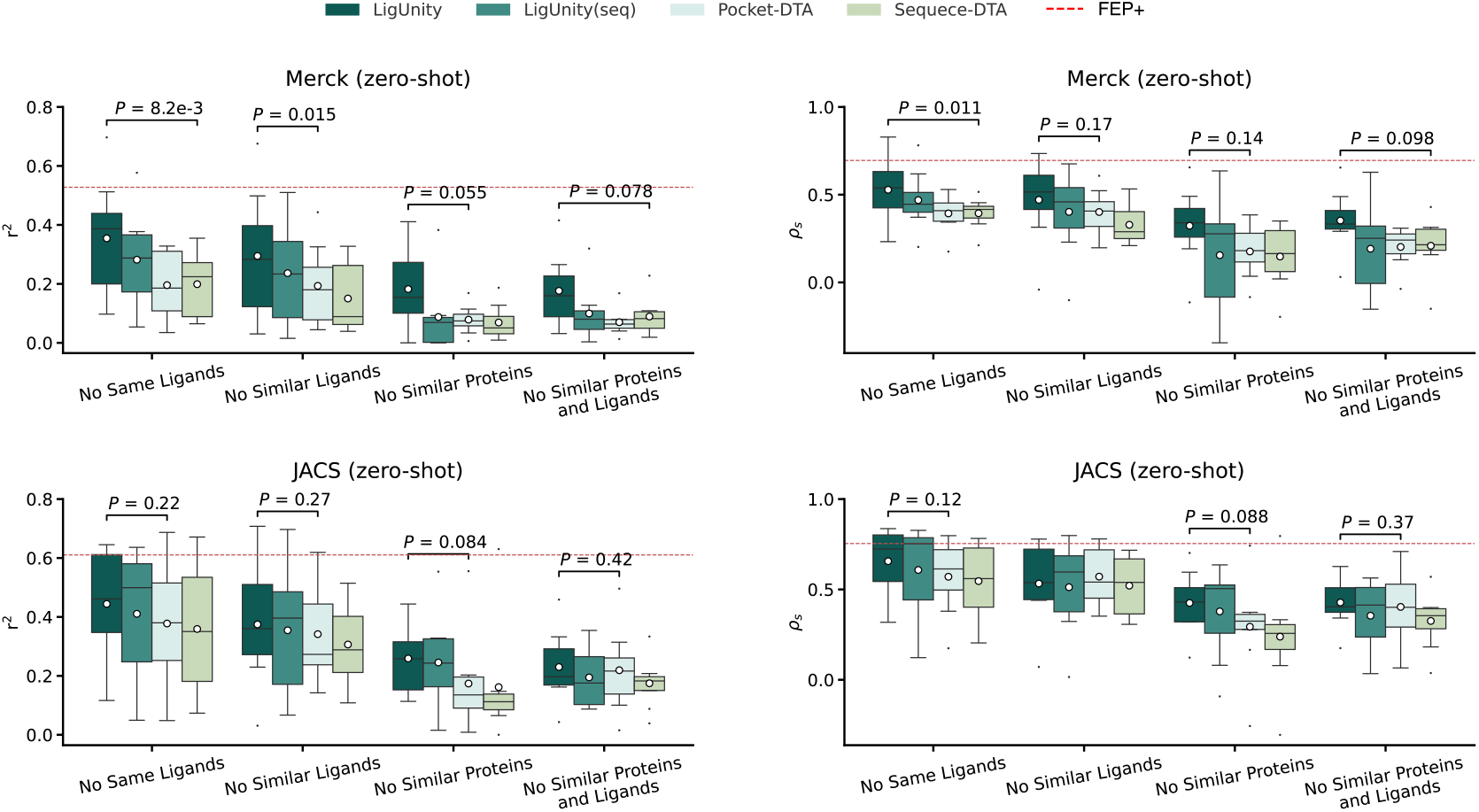
Box plots comparing LigUnity and competing methods across different settings on Merck and JACS benchmarks in terms of *r*^2^ and Spearman’s rank correlation (*ρ*_*s*_). The mean values (white dots) are calculated across n = 8 targets.

**Supplementary Figure 15:**
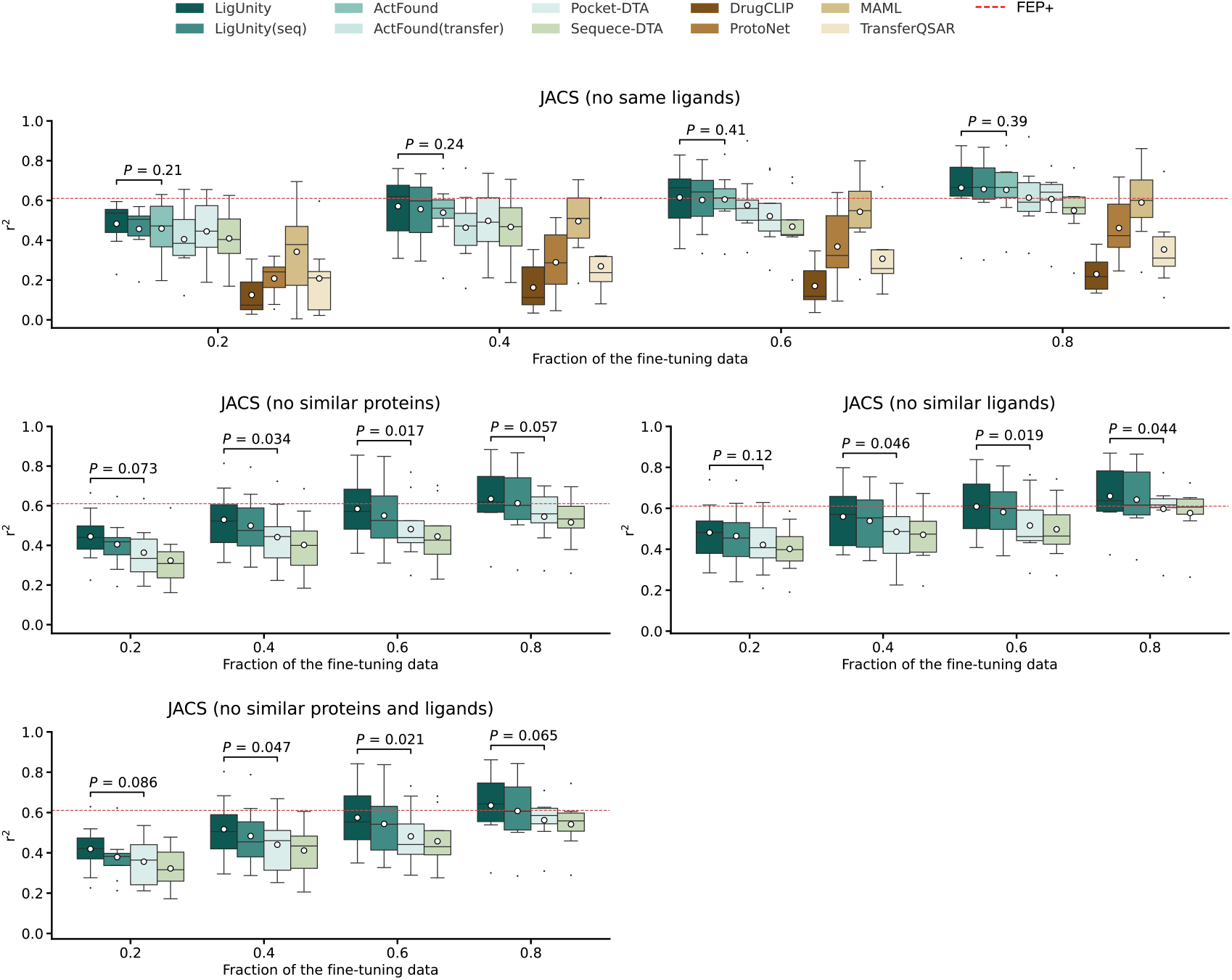
Box plots comparing LigUnity and competing methods across different settings on the JACS benchmark in terms of *r*^2^ and Spearman’s rank correlation (*ρ*_*s*_) when 20%, 40%, 60%, 80% of the experimental binding affinities are used for fine-tuning. The mean values (white dots) are calculated across n = 8 targets.

**Supplementary Figure 16:**
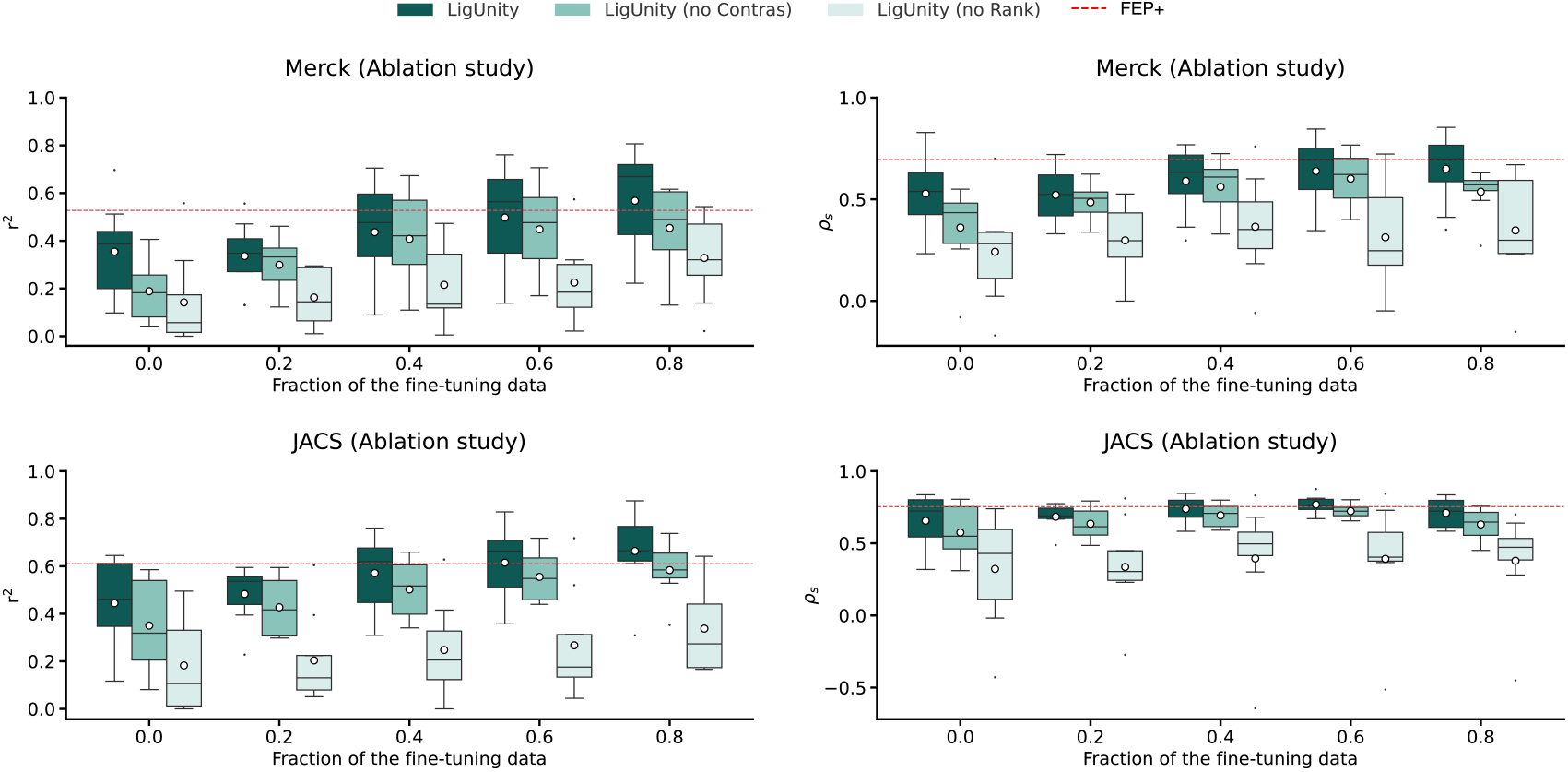
Box plots of ablation study evaluating the contribution of different modules on Merck and JACS benchmarks in terms of *r*^2^ and Spearman’s rank correlation (*ρ*_*s*_) when 0% (zero-shot), 20%, 40%, 60%, 80% of the experimental binding affinities are used for fine-tuning, where “no Contras” means that the contrastive loss for scaffold discrimination are ablated and “no Rank” means that the ranking loss for pharmacophore ranking are ablated. The mean values (white dots) are calculated across n = 8 targets.

**Supplementary Figure 17:**
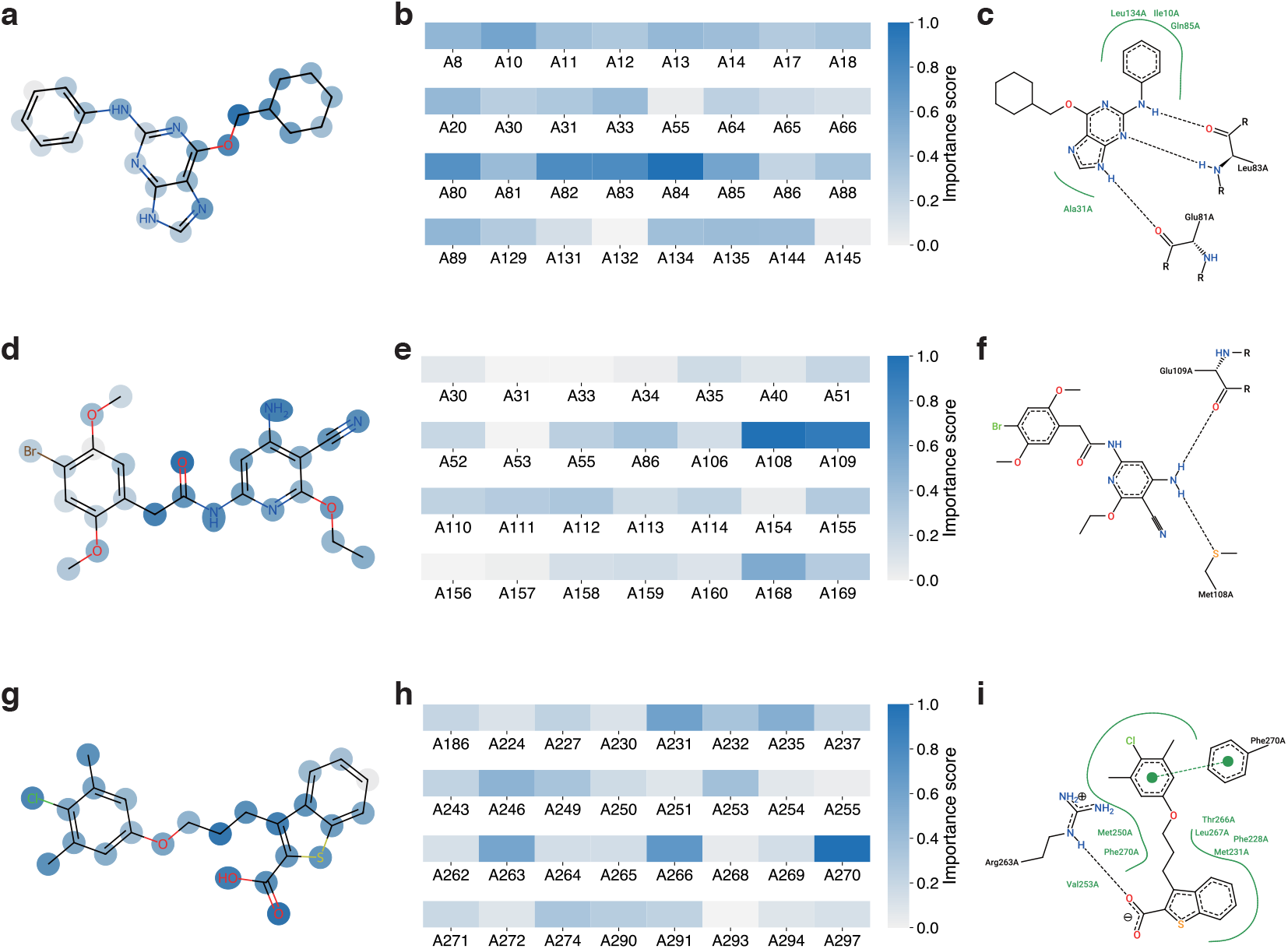
Case study on the CDK2 target **(a**,**b**,**c)**, JNK1 target **(d**,**e**,**f)**, MCL1 **(g, h, i)** target showing the importance score of each ligand atom **(a**,**d**,**g**) and pocket residue **(b**,**e**,**h)** predicted by LigUnity, and 2D interaction graph **(c**,**f**,**i)** showing the non-covalent interaction between the ligand and pocket.

**Supplementary Figure 18:**
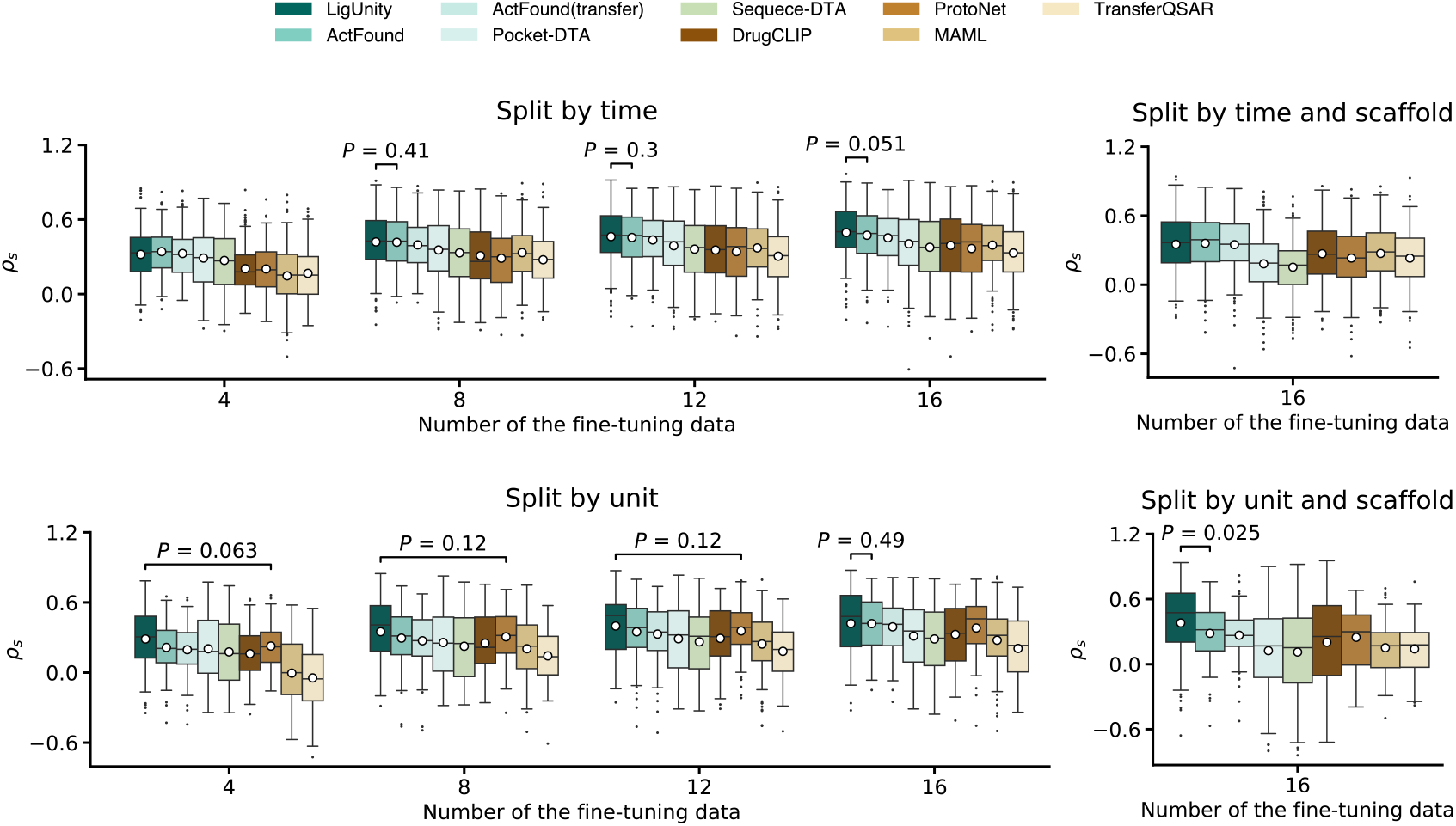
Box plots comparing the binding affinity prediction on the split-by-time and split-by-unit setting in terms of *ρ*_*s*_ when 4, 8, 12, and 16 of the experimental binding affinity is used for fine-tuning. The mean values (white dots) are calculated across n = 161 assays for split-by-time setting and n = 65 assays for split-by-time setting.

**Supplementary Figure 19:**
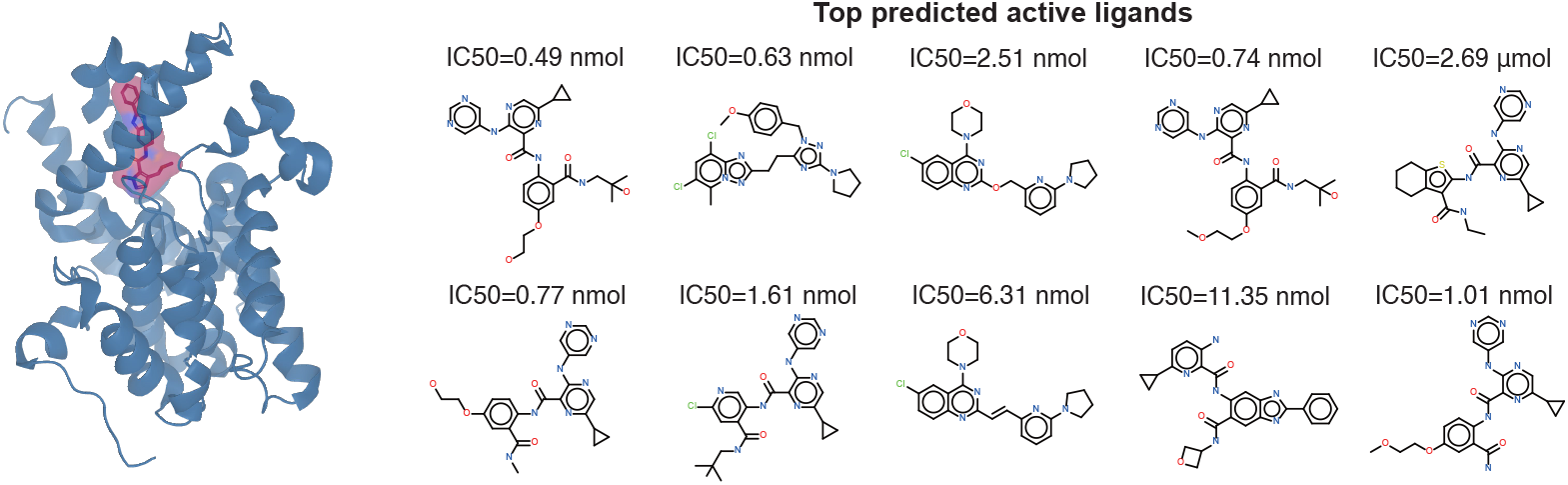
3D structure for the PDE10A target and its active ligand, and the top 10 ligands ranked by affinities predicted using LigUnity, along with their experimental IC50 values.

**Supplementary Figure 20:**
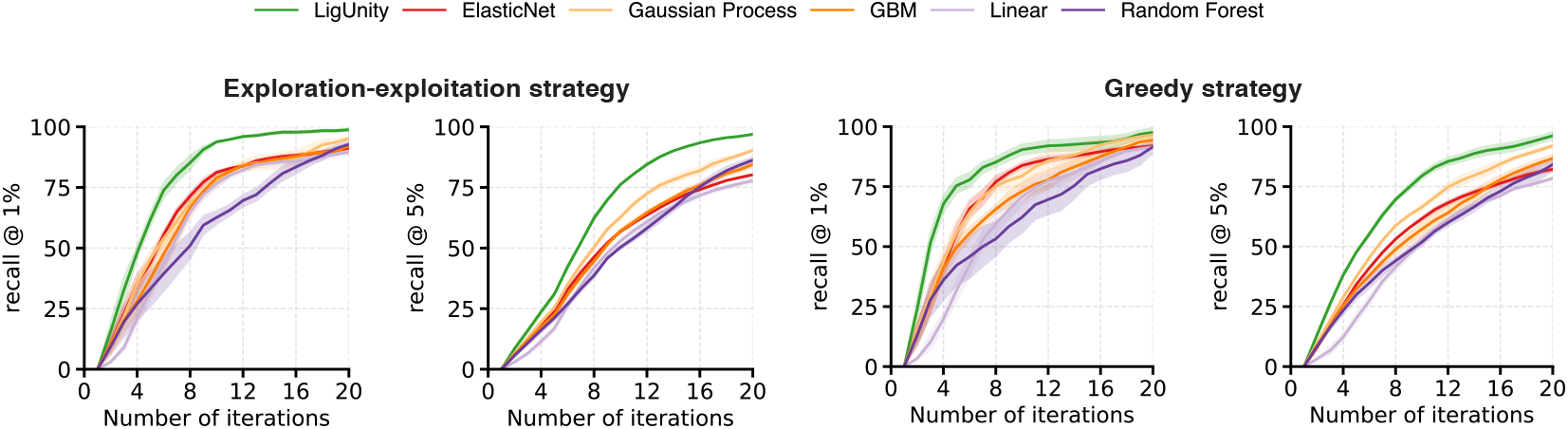
Plots comparing LigUnity and competing methods on the TYK2 dataset in terms of top 1% recall and top 5% recall when using the exploration-exploitation strategy (left) and greedy selection strategy (right).

**Supplementary Figure 21:**
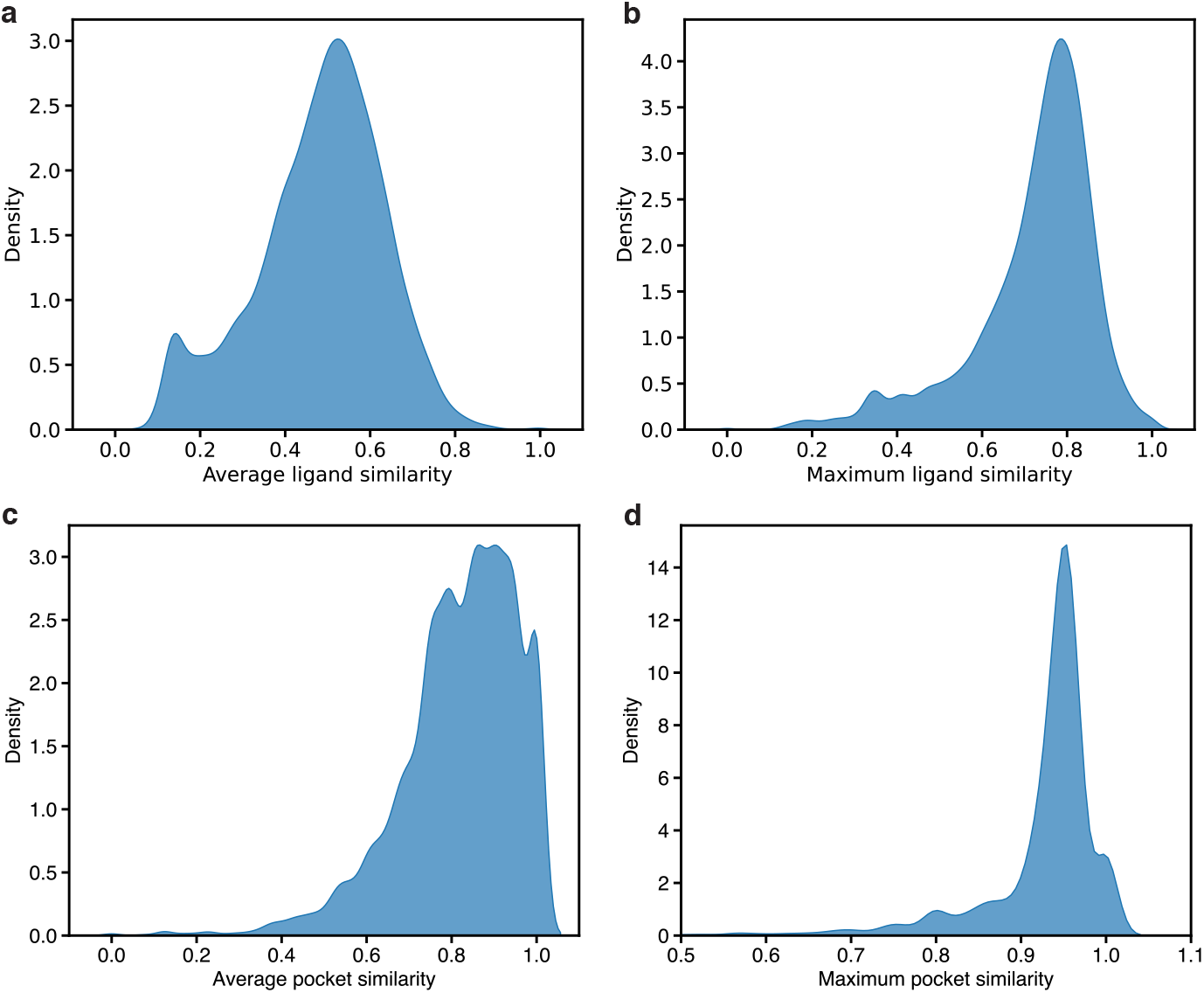
**(a**,) Kernel density estimate (KDE) plot showing average pairwise similarity between any two ligands in an assay. **(b**,) KDE plot showing average similarity between each ligand and its most similar counterpart (nearest neighbor) in an assay. **(c**,) KDE plot showing average pairwise similarity between any two pockets in an assay. **(d**,) KDE plot showing average similarity between each pocket and its most similar counterpart (nearest neighbor) in an assay. The ligand similarity is computed using tanimoto similarity (ECFP4) and the pocket similarity is computed using sequence similarity.

**Supplementary Table 1:** Performance comparison between LigUnity and BIND^78^ on virtual screening benchmarks (DUD-E, LIT-PCBA, and DEKOIS 2.0) in terms of EF 1%, AUROC, and BEDROC score (*α* = 80.5), evaluated under sequence similarity exclusion (90% similarity to test data).

| Method | Metric | DUD-E | LIT-PCBA | DEKOIS 2.0 |
| --- | --- | --- | --- | --- |
| BIND | EF1% | 26.390 (2.008) | 4.450 (1.170) | 15.190 (1.060) |
|  | AUROC | 0.826 (0.018) | 0.564 (0.030) | 0.832 (0.018) |
|  | BEDROC | 0.430 (0.030) | 0.047 (0.010) | 0.453 (0.029) |
| LigUnity | EF1% | 44.781 (2.243) | 4.685 (1.402) | 23.944 (0.992) |
|  | AUROC | 0.871 (0.016) | 0.554 (0.015) | 0.882 (0.014) |
|  | BEDROC | 0.670 (0.032) | 0.060 (0.015) | 0.698 (0.029) |

**Supplementary Table 2:** Performance comparison between LigUnity and PBCNet^15^ on hit-to-lead optimization tasks in terms of Pearson correlation using Merck and JACS benchmarks. Results are shown for zero-shot (0), 6-shot, and 10-shot settings, reporting Pearson’s correlation coefficients. LigUnity is trained following PBCNet’s setting which excludes training assays (congeneric series in PBCNet) identical to assays in FEP benchmarks. We consider two assays identical if they share the same protein and ligands.

| Method | Merck |  |  | JACS |  |  |
| --- | --- | --- | --- | --- | --- | --- |
|  | 0 | 6 | 10 | 0 | 6 | 10 |
| PBCNet | 0.468 | 0.592 | 0.646 | 0.648 | 0.729 | 0.731 |
| LigUnity | 0.494 | 0.631 | 0.676 | 0.690 | 0.746 | 0.775 |

**Supplementary Table 3:** Validation of LigUnity’s ability to learn protein-ligand interaction patterns through targeted residue masking experiments. Performance metrics (square of Pearson correlation *r*^2^ and Spearman correlation *ρ* with standard deviations) are shown for Merck and JACS benchmarks under three conditions: (1) masking key interaction residues identified via ProteinPlus^34^, (2) masking randomly selected residues, and (3) no masking.

| Method | Merck |  | JACS |  |
| --- | --- | --- | --- | --- |
| | $r^2$ | $\rho_s$ | $r^2$ | $\rho_s$ |
| Mask key residue | 0.323 (0.070) | 0.484 (0.066) | 0.354 (0.073) | 0.522 (0.088) |
| Mask random residue | 0.328 (0.067) | 0.497 (0.062) | 0.391 (0.063) | 0.575 (0.074) |
| No mask | 0.333 (0.068) | 0.499 (0.063) | 0.389 (0.064) | 0.578 (0.074) |

**Supplementary Table 4:** Performance comparison of LigUnity under different data curation strategies on virtual screening benchmarks in terms of EF 1%, AUROC, and BEDROC score (*α* = 80.5). “LigUnity (strict ligand)” removes assays with multiple pockets (*>* 10 Å apart), and “LigUnity (strict pocket)” applies a 100% Tanimoto similarity threshold for pocket assignment.

| Benchmark | Metric | LigUnity | LigUnity (strict ligand) | LigUnity (strict pocket) |
| --- | --- | --- | --- | --- |
| DUDE | BEDROC | 0.789 (0.023) | 0.725 (0.026) | 0.732 (0.028) |
|  | AUROC | 0.931 (0.009) | 0.903 (0.013) | 0.905 (0.011) |
|  | EF1 | 52.042 (1.688) | 47.590 (1.865) | 48.374 (2.013) |
| LIT-PCBA | BEDROC | 0.089 (0.027) | 0.082 (0.026) | 0.082 (0.021) |
|  | AUROC | 0.589 (0.029) | 0.553 (0.029) | 0.570 (0.025) |
|  | EF1 | 7.359 (2.488) | 6.803 (2.626) | 7.320 (2.276) |
| DEKOIS 2.0 | BEDROC | 0.849 (0.021) | 0.776 (0.026) | 0.778 (0.025) |
|  | AUROC | 0.941 (0.011) | 0.916 (0.012) | 0.910 (0.011) |
|  | EF1 | 28.212 (0.546) | 25.995 (0.863) | 26.205 (0.785) |

## References

1. Du, X. et al. Insights into protein–ligand interactions: mechanisms, models, and methods. Int. journal molecular sciences 17, 144 (2016).

2. Gorgulla, C. et al. An open-source drug discovery platform enables ultra-large virtual screens. Nature 580, 663–668 (2020).

3. Gentile, F. et al. Artificial intelligence–enabled virtual screening of ultra-large chemical libraries with deep docking. Nat. Protoc. 17, 672–697 (2022).

4. Zhou, G. et al. An artificial intelligence accelerated virtual screening platform for drug discovery. Nat. Commun. 15, 7761 (2024).

5. Heifetz, A., Southey, M., Morao, I., Townsend-Nicholson, A. & Bodkin, M. J. Computational methods used in hit-to-lead and lead optimization stages of structure-based drug discovery. Comput. Methods for GPCR Drug Discov. 375–394 (2018).

6. Huang, L. et al. A dual diffusion model enables 3d molecule generation and lead optimization based on target pockets. Nat. Commun. 15, 2657 (2024).

7. Cerqueira, N. M. et al. Receptor-based virtual screening protocol for drug discovery. Arch. biochemistry biophysics 582, 56–67 (2015).

8. Pagadala, N. S., Syed, K. & Tuszynski, J. A. Software for molecular docking: a review. Biophys. Rev. 9, 91 –102 (2017).

9. Wang, L. et al. Accurate and reliable prediction of relative ligand binding potency in prospective drug discovery by way of a modern free-energy calculation protocol and force field. J. Am. Chem. Soc. 137, 2695–2703 (2015).

10. Wang, E. et al. End-point binding free energy calculation with mm/pbsa and mm/gbsa: strategies and applications in drug design. Chem. reviews 119, 9478–9508 (2019).

11. Cournia, Z., Allen, B. K. & Sherman, W. Relative binding free energy calculations in drug discovery: Recent advances and practical considerations. J. chemical information modeling 57 12, 2911–2937 (2017).

12. Wang, E. et al. End-point binding free energy calculation with mm/pbsa and mm/gbsa: Strategies and applications in drug design. Chem. reviews (2019).

13. Gao, B. et al. DrugCLIP: Contrastive protein-molecule representation learning for virtual screening. Adv. Neural Inf. Process. Syst. (2023).

14. Feng, B. et al. A bioactivity foundation model using pairwise meta-learning. Nat. Mach. Intell. DOI: 10.1038/s42256-024-00876-w (2024).

15. Yu, J. et al. Computing the relative binding affinity of ligands based on a pairwise binding comparison network. Nat. Comput. Sci. 3, 860–872 (2023).

16. Shen, C. et al. A generalized protein–ligand scoring framework with balanced scoring, docking, ranking and screening powers. Chem. Sci. 14, 8129–8146 (2023).

17. Moon, S., Hwang, S.-Y., Lim, J. & Kim, W. Y. Pignet2: a versatile deep learning-based protein–ligand interaction prediction model for binding affinity scoring and virtual screening. Digit. Discov. 3, 287–299 (2024).

18. Wang, Z. et al. A new paradigm for applying deep learning to protein–ligand interaction prediction. Briefings bioinformatics 25, bbae145 (2024).

19. Cao, D. et al. Generic protein–ligand interaction scoring by integrating physical prior knowledge and data augmentation modelling. Nat. Mach. Intell. 1–13 (2024).

20. Gilson, M. K. et al. Bindingdb in 2015: a public database for medicinal chemistry, computational chemistry and systems pharmacology. Nucleic acids research 44, D1045–D1053 (2016).

21. Zdrazil, B. et al. The ChEMBL Database in 2023: a drug discovery platform spanning multiple bioactivity data types and time periods. Nucleic acids research 52, D1180–D1192 (2024).

22. Mysinger, M. M., Carchia, M., Irwin, J. J. & Shoichet, B. K. Directory of useful decoys, enhanced (dud-e): Better ligands and decoys for better benchmarking. J. Medicinal Chem. 55, 6582–6594 (2012).

23. Vogel, S. M., Bauer, M. R. & Boeckler, F. M. Dekois: Demanding evaluation kits for objective in silico screening–a versatile tool for benchmarking docking programs and scoring functions. J. chemical information modeling 51, 2650–2665 (2011).

24. Tran-Nguyen, V.-K., Jacquemard, C. & Rognan, D. Lit-pcba: an unbiased data set for machine learning and virtual screening. J. chemical information modeling 60, 4263–4273 (2020).

25. Friesner, R. A. et al. Glide: a new approach for rapid, accurate docking and scoring. 1. method and assessment of docking accuracy. J. medicinal chemistry 47, 1739–1749 (2004).

26. Trott, O. & Olson, A. J. Autodock vina: improving the speed and accuracy of docking with a new scoring function, efficient optimization, and multithreading. J. computational chemistry 31, 455–461 (2010).

27. Shen, C. et al. Boosting protein–ligand binding pose prediction and virtual screening based on residue–atom distance likelihood potential and graph transformer. J. Medicinal Chem. 65, 10691–10706 (2022).

28. Krasoulis, A., Antonopoulos, N., Pitsikalis, V. & Theodorakis, S. Denvis: scalable and high-throughput virtual screening using graph neural networks with atomic and surface protein pocket features. J. Chem. Inf. Model. 62, 4642–4659 (2022).

29. Öztürk, H., Özgür, A. & Ozkirimli, E. DeepDTA: deep drug–target binding affinity prediction. Bioinformatics 34, i821–i829 (2018).

30. Irwin, J. J. et al. Zinc20—a free ultralarge-scale chemical database for ligand discovery. J. chemical information modeling 60, 6065–6073 (2020).

31. Schindler, C. E. et al. Large-scale assessment of binding free energy calculations in active drug discovery projects. J. Chem. Inf. Model. 60, 5457–5474 (2020).

32. Lyne, P. D., Lamb, M. L. & Saeh, J. C. Accurate prediction of the relative potencies of members of a series of kinase inhibitors using molecular docking and mm-gbsa scoring. J. medicinal chemistry 49, 4805–4808 (2006).

33. Abramson, J. et al. Accurate structure prediction of biomolecular interactions with alphafold 3. Nature 630, 493 (2024).

34. Schöning-Stierand, K. et al. Proteins plus: interactive analysis of protein–ligand binding interfaces. Nucleic acids research 48, W48–W53 (2020).

35. Tosstorff, A. et al. A high quality, industrial data set for binding affinity prediction: performance comparison in different early drug discovery scenarios. J. Comput. Mol. Des. 36, 753–765 (2022).

36. Thompson, J. et al. Optimizing active learning for free energy calculations. Artif. Intell. Life Sci. 2, 100050 (2022).

37. Reker, D. & Schneider, G. Active-learning strategies in computer-assisted drug discovery. Drug discovery today 20, 458–465 (2015).

38. Huang, K. et al. Deeppurpose: a deep learning library for drug–target interaction prediction. Bioinformatics 36, 5545–5547 (2020).

39. Chatterjee, A. et al. Improving the generalizability of protein-ligand binding predictions with ai-bind. Nat. communications 14, 1989 (2023).

40. Luo, Y., Liu, Y. & Peng, J. Calibrated geometric deep learning improves kinase–drug binding predictions. Nat. Mach. Intell. 5, 1390–1401 (2023).

41. Pei, Q. et al. Breaking the barriers of data scarcity in drug–target affinity prediction. Briefings Bioinforma. 24, bbad386 (2023).

42. Nguyen, T. et al. GraphDTA: predicting drug–target binding affinity with graph neural networks. Bioinformatics 37, 1140–1147 (2021).

43. Zhao, Q., Xiao, F., Yang, M., Li, Y. & Wang, J. AttentionDTA: prediction of drug–target binding affinity using attention model. In 2019 IEEE international conference on bioinformatics and biomedicine (BIBM), 64–69 (IEEE, 2019).

44. Yang, Z., Zhong, W., Zhao, L. & Chen, C. Y.-C. MGraphDTA: deep multiscale graph neural network for explainable drug–target binding affinity prediction. Chem. science 13, 816–833 (2022).

45. Lin, S., Shi, C. & Chen, J. Generalizeddta: combining pre-training and multi-task learning to predict drug-target binding affinity for unknown drug discovery. BMC bioinformatics 23, 367 (2022).

46. Yuan, W., Chen, G. & Chen, C. Y.-C. FusionDTA: attention-based feature polymerizer and knowledge distillation for drug-target binding affinity prediction. Briefings Bioinforma. 23, bbab506 (2022).

47. Zhang, X. et al. Efficient and accurate large library ligand docking with karmadock. Nat. Comput. Sci. 3, 789–804 (2023).

48. Durrant, J. D. & McCammon, J. A. Nnscore 2.0: a neural-network receptor–ligand scoring function. J. chemical information modeling 51, 2897–2903 (2011).

49. Ballester, P. J. & Mitchell, J. B. A machine learning approach to predicting protein–ligand binding affinity with applications to molecular docking. Bioinformatics 26, 1169–1175 (2010).

50. Wang, C. & Zhang, Y. Improving scoring-docking-screening powers of protein–ligand scoring functions using random forest. J. computational chemistry 38, 169–177 (2017).

51. Lu, J., Hou, X., Wang, C. & Zhang, Y. Incorporating explicit water molecules and ligand conformation stability in machine-learning scoring functions. J. chemical information modeling 59, 4540–4549 (2019).

52. Yang, C. & Zhang, Y. Delta machine learning to improve scoring-ranking-screening performances of protein–ligand scoring functions. J. Chem. Inf. Model. 62, 2696–2712 (2022).

53. Stepniewska-Dziubinska, M. M., Zielenkiewicz, P. & Siedlecki, P. Pafnucy - a deep neural network for structure-based drug discovery. ArXiv abs/1712.07042 (2017).

54. Zheng, L., Fan, J. & Mu, Y. Onionnet: a multiple-layer intermolecular-contact-based convolutional neural network for protein–ligand binding affinity prediction. ACS omega 4, 15956–15965 (2019).

55. Moon, S., Zhung, W., Yang, S., Lim, J. & Kim, W. Y. Pignet: a physics-informed deep learning model toward generalized drug–target interaction predictions. Chem. Sci. 13, 3661–3673 (2022).

56. Zhang, X. et al. Planet: a multi-objective graph neural network model for protein–ligand binding affinity prediction. J. Chem. Inf. Model. 64, 2205–2220 (2023).

57. McNutt, A. T. et al. Gnina 1.0: molecular docking with deep learning. J. cheminformatics 13, 43 (2021).

58. Kalliokoski, T., Kramer, C., Vulpetti, A. & Gedeck, P. Comparability of mixed ic50 data–a statistical analysis. PloS one 8, e61007 (2013).

59. Zhou, G. et al. Uni-mol: A universal 3d molecular representation learning framework. In The Eleventh International Conference on Learning Representations (2023).

60. Li, H. et al. Decoupled peak property learning for efficient and interpretable electronic circular dichroism spectrum prediction. Nat. Comput. Sci. 1–11 (2025).

61. Cao, Z., Qin, T., Liu, T.-Y., Tsai, M.-F. & Li, H. Learning to rank: from pairwise approach to listwise approach. In International Conference on Machine Learning (2007).

62. Mayr, A. et al. Large-scale comparison of machine learning methods for drug target prediction on chembl. Chem. Sci. 9, 5441–5451 (2018).

63. Ross, G. A. et al. The maximal and current accuracy of rigorous protein-ligand binding free energy calculations. Commun. Chem. 6, 222 (2023).

64. Lin, Z. et al. Evolutionary-scale prediction of atomic-level protein structure with a language model. Science 379, 1123–1130 (2023).

65. Zardecki, C., Dutta, S., Goodsell, D. S., Voigt, M. & Burley, S. K. Rcsb protein data bank: A resource for chemical, biochemical, and structural explorations of large and small biomolecules. J. Chem. Educ. 93, 569–575, DOI: 10.1021/acs.jchemed.5b00404 (2016).

66. Maggiora, G., Vogt, M., Stumpfe, D. & Bajorath, J. Molecular similarity in medicinal chemistry. J. Medicinal Chem. 57, 3186–3204, DOI: 10.1021/jm401411z (2014). PMID: 24151987, https://doi.org/10.1021/jm401411z.

67. Yang, J., Roy, A. & Zhang, Y. Biolip: a semi-manually curated database for biologically relevant ligand–protein interactions. Nucleic acids research 41, D1096–D1103 (2012).

68. Martin, E. J. et al. All-assay-Max2 pQSAR: activity predictions as accurate as four-concentration ic50s for 8558 novartis assays. J. chemical information modeling 59, 4450–4459 (2019).

69. Berman, H. M. et al. The Protein Data Bank. Nucleic acids research 28, 235–242 (2000).

70. Wang, R., Fang, X., Lu, Y. & Wang, S. The pdbbind database: Collection of binding affinities for protein-ligand complexes with known three-dimensional structures. J. Medicinal Chem. 47, 2977–2980 (2004).

71. Huey, R., Morris, G. M., Olson, A. J. & Goodsell, D. S. A semiempirical free energy force field with charge-based desolvation. J. computational chemistry 28, 1145–1152 (2007).

72. Quiroga, R. & Villarreal, M. A. Vinardo: A scoring function based on autodock vina improves scoring, docking, and virtual screening. PloS one 11, e0155183 (2016).

73. Verdonk, M. L., Cole, J. C., Hartshorn, M. J., Murray, C. W. & Taylor, R. D. Improved protein–ligand docking using gold. Proteins: Struct. Funct. Bioinforma. 52, 609–623 (2003).

74. Jain, A. N. Surflex: fully automatic flexible molecular docking using a molecular similarity-based search engine. J. medicinal chemistry 46, 499–511 (2003).

75. Rarey, M., Kramer, B., Lengauer, T. & Klebe, G. A fast flexible docking method using an incremental construction algorithm. J. molecular biology 261, 470–489 (1996).

76. Jiménez, J., Skalic, M., Martinez-Rosell, G. & De Fabritiis, G. K deep: protein–ligand absolute binding affinity prediction via 3d-convolutional neural networks. J. chemical information modeling 58, 287–296 (2018).

77. Lu, W. et al. Tankbind: Trigonometry-aware neural networks for drug-protein binding structure prediction. Adv. neural information processing systems 35, 7236–7249 (2022).

78. Lam, H. Y. I., Guan, J. S., Ong, X. E., Pincket, R. & Mu, Y. Protein language models are performant in structure-free virtual screening. Briefings Bioinforma. 25, bbae480 (2024).

79. Zou, H. & Hastie, T. Regularization and variable selection via the elastic net. J. Royal Stat. Soc. Ser. B: Stat. Methodol. 67, 301–320 (2005).

80. Williams, C. & Rasmussen, C. Gaussian processes for regression. Adv. neural information processing systems 8 (1995).

81. DiFranzo, A., Sheridan, R. P., Liaw, A. & Tudor, M. Nearest neighbor gaussian process for quantitative structure–activity relationships. J. Chem. Inf. Model. 60, 4653–4663 (2020).

82. Sheridan, R. P., Wang, W. M., Liaw, A., Ma, J. & Gifford, E. M. Extreme gradient boosting as a method for quantitative structure–activity relationships. J. chemical information modeling 56, 2353–2360 (2016).

83. Svetnik, V. et al. Random forest: a classification and regression tool for compound classification and qsar modeling. J. chemical information computer sciences 43, 1947–1958 (2003).

84. Lu, C. et al. OPLS4: Improving force field accuracy on challenging regimes of chemical space. J. chemical theory computation 17, 4291–4300 (2021).

85. Rocklin, G. J., Mobley, D. L. & Dill, K. A. Separated topologies—a method for relative binding free energy calculations using orientational restraints. The J. chemical physics 138 (2013).

